# Molecular and functional characterization of the *Drosophila melanogaster* conserved smORFome

**DOI:** 10.1101/2022.04.24.489283

**Authors:** Justin A. Bosch, Nathan Keith, Felipe Escobedo, William W. Fisher, James Thai LaGraff, Jorden Rabasco, Kenneth H. Wan, Richard Weiszmann, Yanhui Hu, Shu Kondo, James B. Brown, Norbert Perrimon, Susan E. Celniker

## Abstract

Short polypeptides encoded by small open reading frames (smORFs) are ubiquitously found in eukaryotic genomes and are important regulators of physiology, development, and mitochondrial processes. Here, we focus on a subset of 298 smORFs that are evolutionarily conserved between *Drosophila melanogaster* and humans. Many of these smORFs are conserved broadly in the bilaterian lineage, with ∼182 conserved in plants. Within these conserved smORFs, we observed remarkably heterogenous spatial and temporal expression patterns – indicating wide-spread tissue-specific and stage-specific mitochondrial architectures. In addition, an analysis of annotated functional domains revealed a predicted enrichment of smORF polypeptides localizing to mitochondria. We conducted an embryonic ribosome profiling experiment finding support for translation of 137 of these smORFs during embryogenesis. We further embarked on functional characterization using CRISPR knockout/activation, RNAi knockdown, and cDNA overexpression, revealing diverse phenotypes. This study underscores the importance of identifying smORF function in disease and phenotypic diversity.

## Introduction

Genome annotations have often overlooked proteins with less than 100 amino acids, although many have been shown to play important roles in development and physiology and are pervasive across the Tree of Life (Couso and Patraquim, 2017; Guerra-Almeida et al., 2021; Plaza et al., 2017). While some small proteins are cleavage products of longer proteins, many others are encoded in the genome by small open reading frames (smORF genes; δ 100 amino acids). Strikingly, it has been reported that human disease-associated variants from genome-wide association studies (GWAS) are enriched in smORF genes (Jain et al., 2023). These estimates underscore the functional roles of smORFs and relevance to human diseases.

Advances in proteomics and next-generation sequencing (NGS) technologies have led to a significantly improved annotation of smORF genes. There is evidence for >2,500 smORF genes in humans (Martinez et al., 2020), and there are over 1,000 annotated smORF genes in the *Drosophila melanogaster* genome (Adams et al., 2000). Some smORF genes are important regulators of physiology, development and metabolism and encode hormones (Pearson et al., 1993), neurotransmitters (Snyder and Innis, 1979), ligands and cofactors (Yang et al., 2015), RNA and DNA binding factors, and components of ribonucleoproteins (Henras et al., 1998). Interestingly, a number of them are involved in numerous mitochondrial functions and processes (Bosch et al., 2022; Rathore et al., 2018; Stein et al., 2018). Notably, studies in insects have defined the functions of several smORF peptides, such as Tarsal-less/mille-pattes/polished-rice (Chanut-Delalande et al., 2014; Galindo et al., 2007; Kondo et al., 2010; Savard et al., 2006), Brd (Bardin and Schweisguth, 2006; Chanet and Schweisguth, 2012; Lai et al., 2000), and Pgc (Hanyu-Nakamura et al., 2008).

Although smORF genes are prevalent in metazoan genomes, a surprisingly small number of these genes are evolutionarily conserved in animals suggesting a high birth and death rate for these genes (Mackowiak et al., 2015; Martinez et al., 2020). For instance, there are over 2,500 smORF sequences with ribosome profiling evidence of translation across human cell lines, but only 273 of these human smORFs in the mouse genome based on computational analysis (Martinez et al., 2020). smORF genes with deep evolutionary conservation are therefore of particular interest because of their assumed importance to the health and fitness across Metazoa and their implications for human health and disease. Again, studies in Drosophila have pioneered the bioinformatic identification of smORF peptides, using techniques such as amino acid conservation, ribosomal profiling, and proteomics (Fabre et al., 2022; Ladoukakis et al., 2011; Patraquim et al., 2022; Zhang et al., 2022).

Here, we characterize a collection of 298 fly smORFs conserved with human. For a subset of these smORFs, we describe their spatial expression patterns and phenotypes associated with gene loss of function or overexpression. Many of the pronounced phenotypes are associated with expression in neural tissue, and genes encoding mitochondrial proteins. In addition, several phenotypes were detectable only in flies subjected to stressful diets. This study serves as a resource for the functional annotation of this diverse and under-studied class of genes.

## Results

### Deep conservation of smORFs

We identified 298 smORF genes that are evolutionarily conserved between humans and *Drosophila melanogaster*, which we refer to as conserved smORFs (Materials and Methods; Supplemental File 1). All 298 are currently annotated as protein coding. We further analyzed conservation of these smORFs with the well-annotated transcriptomes of zebrafish (*Danio rerio*), nematodes (*Caenorhabditis elegans)* and thale cress (*Arabidopsis thaliana)* (Figure 1A). There are 274 conserved in zebrafish, and 239 conserved in *C. elegans*. Notably, 182 conserved smORFs were also conserved in *Arabidopsis*, providing evidence for the functional importance of this dataset in non-Bilateria eukaryotes. Finally, amino acid alignment of many of the human-fly conserved smORFs, such as *bc10*, *CG42497*, and *Tim10,* reveal that they are also conserved among other invertebrate and vertebrate species (Figure 1B).

**Figure 1.**
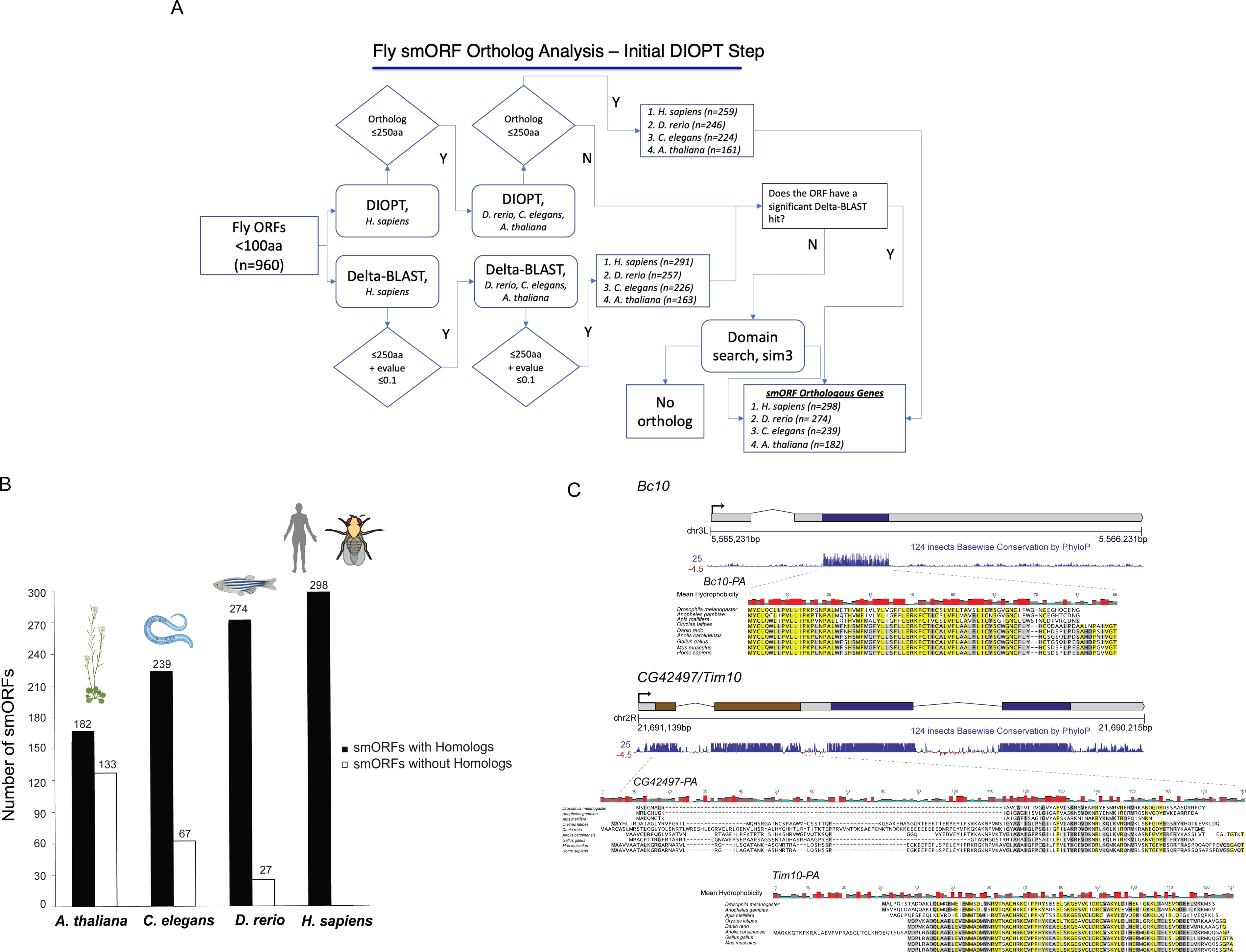
Conservation of smORF dataset. **(*A*)** Flowthrough of bioinformatic identification of 298 fly-human conserved smORFs **(*B*)** Number of conserved smORFs in dataset with and without homologs in a selection of species with well-annotated transcriptomes. ***(C*)** Multiple-species alignments of conserved smORFs, including mean amino acid hydrophobicity at each alignment position. *bc10* (upper transcript) is encodes one smORF, whereas *CG42497* and *Tim10* (lower transcript) is polycistronic.

Of the 298 *Drosophila* conserved smORFs, 32 are polycistronic (Supplemental File 1, see example in Figure 1C). Remarkably, while the individual smORFs that reside in *Drosophila* polycistronic transcripts are evolutionarily conserved, their polycistronic structure is generally not - indicating a complex evolutionary history. Interestingly, there are three conserved smORFs encoded by polycistronic transcripts in both fly and *C. elegans*, *CG42372, CG42375* and *Mocs2A*. Between flies and zebrafish, there are two smORFs encoded by polycistronic transcripts in both species, *CG42497* and *Mocs2A*. However, between flies and humans, or flies and *Arabidopsis,* there are none. All non-fly smORF orthologs are currently annotated as protein-coding.

### Gene Ontology analysis of conserved smORFs

The functions of conserved smORFs are diverse, as with any broad category of genes. Gene Ontology (GO) Cellular Component enrichment analysis of conserved smORFs determined that the majority of significantly enriched GO terms are associated with mitochondrial function and localization (Figure 2, Supplemental File 2). Indeed, 66 conserved smORFs are predicted to be involved in mitochondrial function (Supplemental File 2). “Mitochondrion” (*P* = 1.46 x 10^-51^) which contains 63 conserved smORFs is the most significantly enriched GO Cellular Component terms in this dataset, with “Mitochondrial Envelope” as the second most significantly enriched GO Cellular Component term (*P* = 2.2 x 10^-49^) (Figure 2). Additional significantly enriched terms include mitochondrial inner membrane (*P* = 1.4 x 10^-45^) and cytochrome complex (*P* = 6.09 x 10^-24^) (Figure 2).

**Figure 2.**
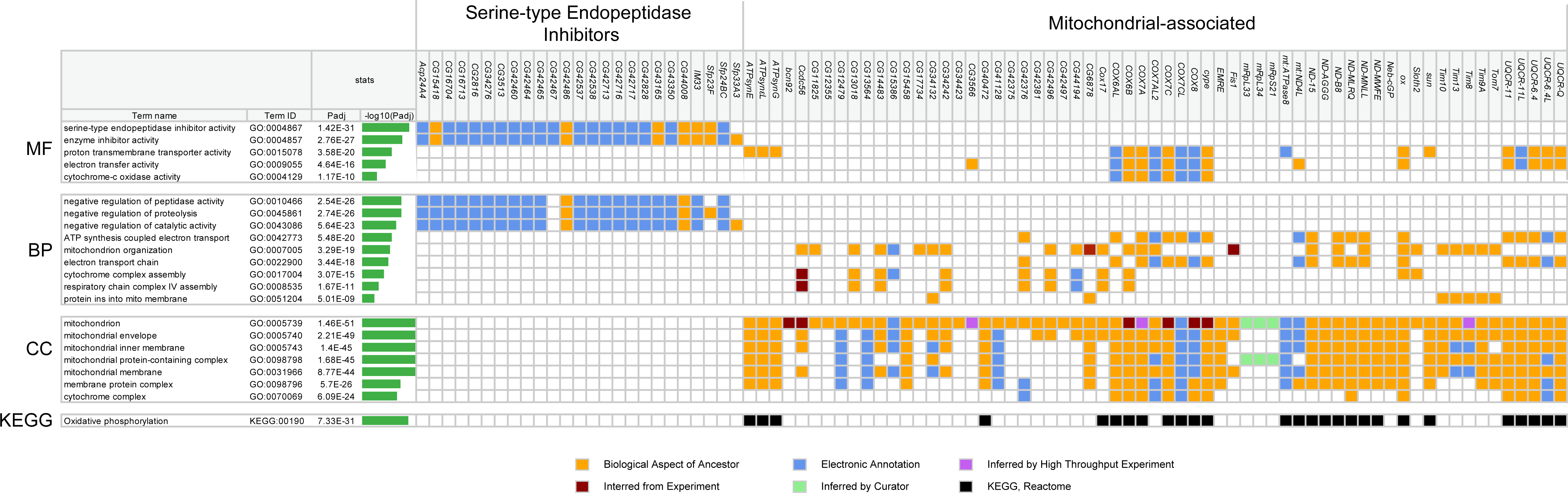
Gene Ontology (GO) and KEGG enrichment analysis of conserved smORFs. Significantly enriched GO terms for molecular function (“MF”), biological process (“BP”), cellular component (“CC”) are plotted. GO and KEGG enrichment analyses were performed with g:Profiler (Raudvere et al., 2019). Significantly enriched terms <10^-5^ are shown that also encompass all conserved smORFs classified as serine-type endopeptidase inhibitors and mitochondria-associated conserved smORFs.

The oxidative phosphorylation pathway is the only significantly enriched pathway in the smORF dataset (*P* = 7.33 x 10^-31^; (Kanehisa et al., 2021) KEGG; Supplemental File 2, Figure 2, Figure S1). Four of the genes in the oxidative phosphorylation pathway, *COX6CL*, *CG40472, COX7CL,* and *UQCR-6.4L,* are paralogs of annotated fly genes: *cyclope* (*cype*) encoding the *cytochrome c oxidase subunit 6C*, *NADH dehydrogenase (ubiquinone) AGGG subunit* (*ND-AGGG*)*, Ubiquinol-cytochrome c reductase 6.4 kDa subunit* (*UQCR-6.4*) and *Cytochrome c oxidase subunit 7C* (*COX7C*). Interestingly, *COX6CL, COX7CL and UQCR-6.4L* are primarily expressed in the adult testis, whereas their paralogs (*cype*, *COX7C,* and *UQCR-6.4*, respectively) are ubiquitously expressed (Brown et al., 2014).

In contrast to the GO Cellular Component enrichment analysis, the most significantly enriched Biological Function and Molecular Process GO terms are related to serine-endopeptidase inhibitor activity (Figure 2; Supplemental File 2). For instance, the most significantly enriched Molecular Function GO term is serine-type endopeptidase inhibitor activity (*P* = 2.91 x 10^-22^) (Figure 2). Notably, 32 of the 80 predicted serine-endopeptidase inhibitors in the *D. melanogaster* genome are in the conserved smORF dataset. Of these 32 smORFs, 22 contain a predicted pancreatic trypsin inhibitor Kunitz domain (Interpro: IPR036880), and 10 smORFs contain a predicted a Kazal domain (Interpro: IPR036058) - providing evidence that these 32 conserved smORFs are serine-endopeptidase inhibitors (Rawlings et al., 2004). Additionally, all 32 of these predicted serine-endopeptidase inhibitors contain an N-terminal secretion signal (SignalP6.0; 0.982 or higher score) (Teufel et al., 2022).

### In situ imaging of smORF mRNA expression in embryos

To better understand the functional roles of smORFs in *Drosophila*, we performed in situ mRNA hybridization to visualize smORF expression during embryogenesis (Supplemental File 3; https://insitu.fruitfly.org/). Of these, organ-specific expression (i.e., “patterned” expression) could be assigned for 143 conserved smORFs during embryonic development (Supplemental File 3).

In addition to patterned expression, conserved smORFs can also be classified as being “maternally deposited”, and therefore ubiquitously expressed in the embryo in the earliest developmental stage(s), and/or classified as being “ubiquitously expressed”, where expression is observed throughout the full embryo after the earliest developmental stage (Supplemental File 4). We found that 59 conserved smORFs were classified as only being maternally deposited and/or being ubiquitously expressed (i.e., expressed throughout the entire embryo without assigned organ patterns) in at least one embryonic stage. Of these 59 smORFs, *CG15456* and *UQCR-6.4L* are the only smORF with no observed expression after maternal deposition (i.e., after initial embryonic stages 1-3).

The remaining 86 smORFs revealed no observed expression in the embryo. In each case this was concordant with RNA-seq (Brown et al., 2018): these smORFs are expressed later in development, and in specific tissues, or under stress conditions (Brown et al., 2018).

### In situ imaging of mitochondria-associated smORF mRNA expression

Notably, mitochondria-associated conserved smORFs exhibited heterogenous spatial expression patterns, indicating tissue- and stage-specific mitochondrial architectures, and, by extension, tissue- and stage-specific mitochondrial functions (Figure 3). Of these, expression could be assigned as patterned for 42 smORFs. (Figure 3). We clustered these mitochondria-associated conserved smORFs with patterned expression into six groups - showing extensive inter- and intra-group heterogeneity in organ system expression patterns (Figure 3).

**Figure 3.**
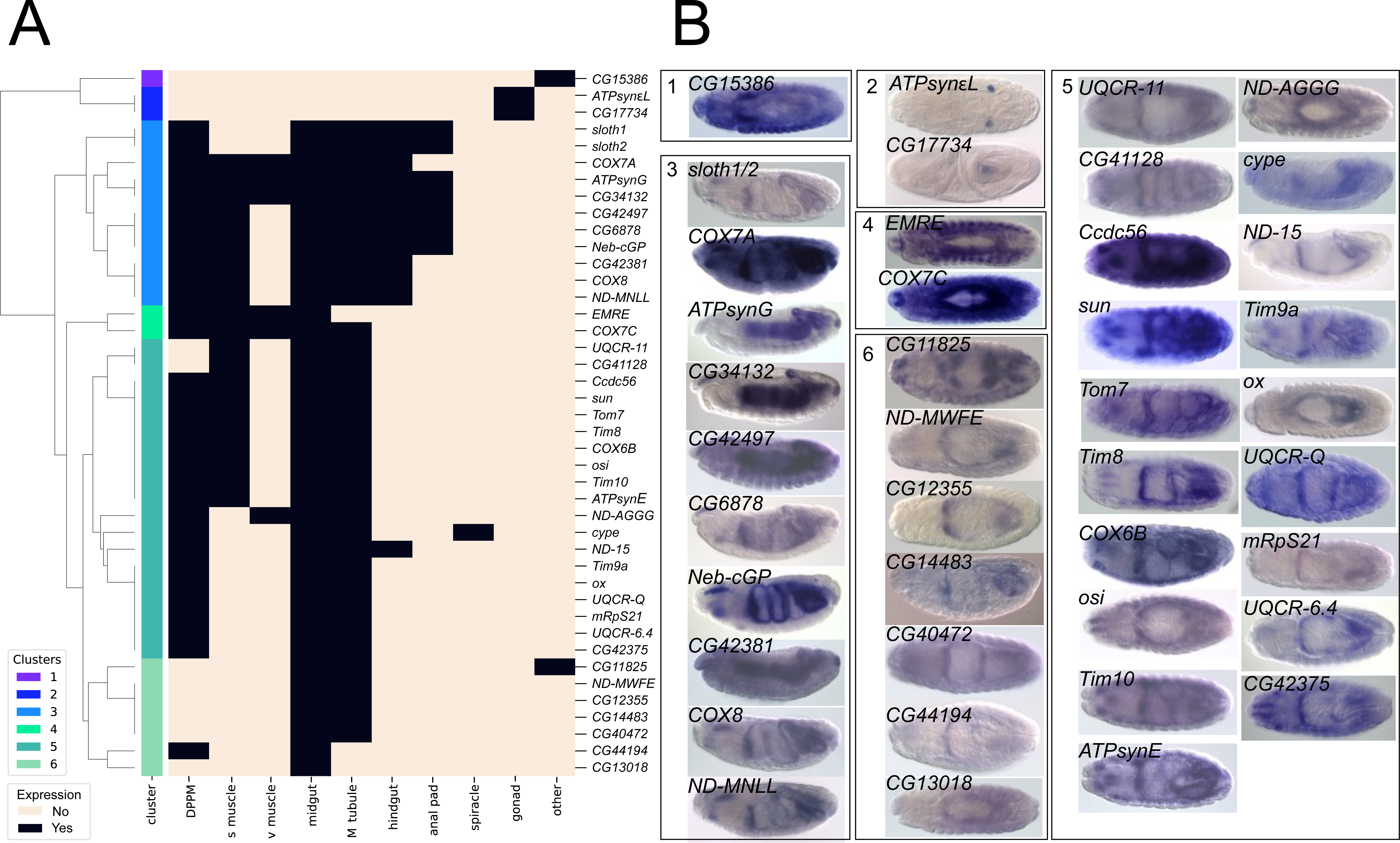
*In situ* mRNA hybridization patterns of mitochondria-associated smORFs. **(*A*)** Clustering of mitochondria-associated conserved smORF *in situ* mRNA expression patterns (Weiszmann et al., 2009). For each mitochondrial conserved smORF, the organs where expression patterns were assigned are represented by red boxes, while blue boxes represent no annotated expression. Expression patterns across embryo stages are collapsed. “DPPM” = dorsal prothoracic pharyngeal muscle; “MT” = Malpighian tubules; “VNC” = ventral nerve cord; “D and V epidermis” = dorsal and ventral epidermis. **(*B*)** In-situ mRNA hybridization images for each mitochondria-associated conserved smORF with patterned expression. Each image was taken between embryonic stages 13-16.

Nearly all (42/45) patterned mitochondria-associated conserved smORFs were maternally deposited (Supplemental Files 3 and 4), and 32/34 were annotated as being ubiquitously expressed in at least one later embryonic stage (Supplementary File 3). Of the two conserved smORFs without ubiquitous expression after maternal deposition, one (*Cox17*) shows robust embryonic expression under RNA-seq (Brown et al., 2014), but *in situ* hybridization was unsuccessful; while the other is only annotated as patterned post maternal deposition (*CG17734*) (Supplemental File 4). However, nearly all patterned, mitochondrial smORFs (33/34) exhibit complex and tissue specific expression patterns after early embryonic ubiquitous expression (usually due to maternal deposition) (Supplemental File 4). Hence, the zygotic regulation of structural and functional components of mitochondria is highly patterned and specific to individual tissues and cell types.

### Ribosome profiling and proteomics analysis to support the translation of conserved smORFs

To verify the translation of conserved smORFs, we performed ribosome profiling for six, 2-hour embryonic stages and six, 0-24 hr. mixed stage embryo samples (Figure 4; Figure S2, Supplemental Files 5 and 6). We find evidence of translation for 137 (46%) conserved smORFs from these embryonic samples. We find that 42 (14%) conserved smORFs are detected in only a single embryonic stage (Supplemental File 5), consistent with transcriptional evidence from extensive RNA-seq experiments in a developmental time course (Brown et al., 2014; Graveley et al., 2011).

**Figure 4.**
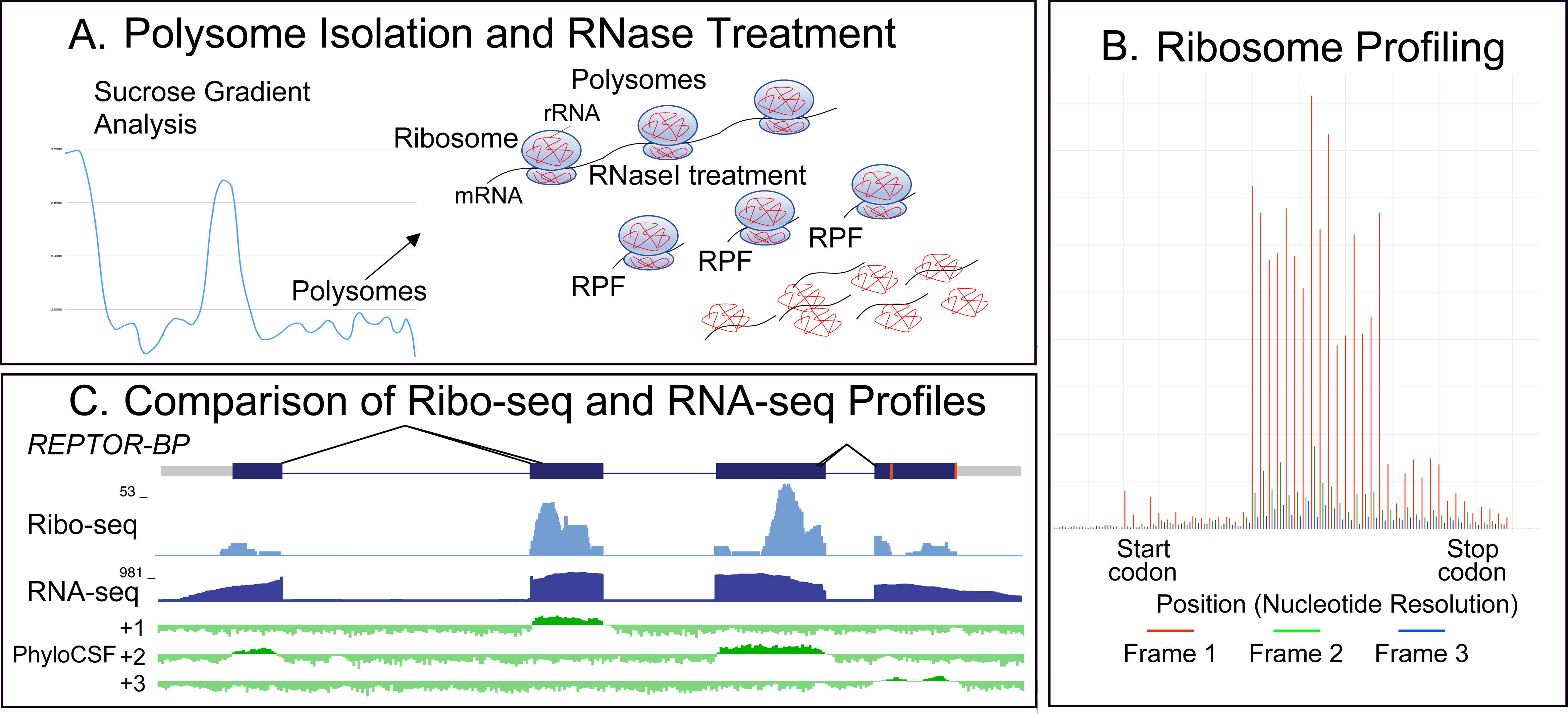
Ribosome Profiling. **(*A*)** Overview of ribosome profiling workflow, where polysomes are isolated followed by digestion of inter-ribosome RNA. Ribosome protected fragments are then collected. **(*B*)** After sequencing, the number of in frame reads are analyzed to determine if RPFs were successfully sequenced. Distribution of tags per million (TPM) for six, embryonic time periods. **(*C*)** Comparison of ribosome profiling sequencing to mRNA-seq sequencing showing ribosome profiling libraries are constrained to CDS while mRNA libraries map to the entire annotated transcript.

We analyzed published *Drosophila* proteomics datasets (Aradska et al., 2015; Brown et al., 2018; Bucio-Mendez et al., 2019; Cammarato et al., 2011; Casas-Vila et al., 2017; Chen et al., 2015; Dorus et al., 2006; Fabre et al., 2022; Wasbrough et al., 2010) that provided translation and/or polypeptide support for 186 total conserved smORFs (Supplemental File 5). Our polysome experiments provide support for an additional 22 smORFs, leaving 90 with no proteomics support. Analysis of comprehensive *Drosophila* modENCODE mRNA expression data (Brown et al., 2014) revealed 53/90 (59%) of the conserved smORFs without evidence for translation or peptides are maximally expressed in testes or accessory gland (Figure S3). Notably, 47/53 (89%) of these conserved smORFs that are maximally expressed in testes and accessory gland are classified as having no or low mRNA expression throughout embryogenesis, and their expression is testes- and/or accessory gland-specific (Brown et al., 2014; Graveley et al., 2011). Interestingly, 40/47 (85%) of these smORFs have a human homolog that are expressed in testes (GTEx Portal, TPM >1).

### Functional analysis of smORF genes by F1 CRISPR screens

Next, we assessed if conserved smORFs have important biological functions in *Drosophila* by modifying their gene function in vivo. First, to knock out (KO) smORF gene function in a systematic manner, we used a CRISPR/Cas9-based transgenic crossing strategy (Port et al., 2020; Zirin et al., 2020) where a Cas9-expressing line is crossed with sgRNA lines that target 5’ coding sequence (sgRNA-KO). The resulting progeny will contain somatic indels in the target gene that disrupt gene function. We generated a collection of 177 sgRNA-KO lines that target 165 smORF genes (Supplemental File 7). Each smORF sgRNA-KO line was crossed with *Act5c-Cas9* (ubiquitous Cas9) and F1 progeny were screened for defects in viability, morphology, or gross motor behavior (Figure 5A). Of the 115 sgRNA-KO lines tested, 14 (representing 14 genes) gave no mutant adult progeny or very few compared to controls (Figure 5B; Supplemental File 7), suggesting that they are essential genes. No other obvious morphological or behavioral phenotypes were observed for the remaining sgRNA-KO lines.

**Figure 5.**
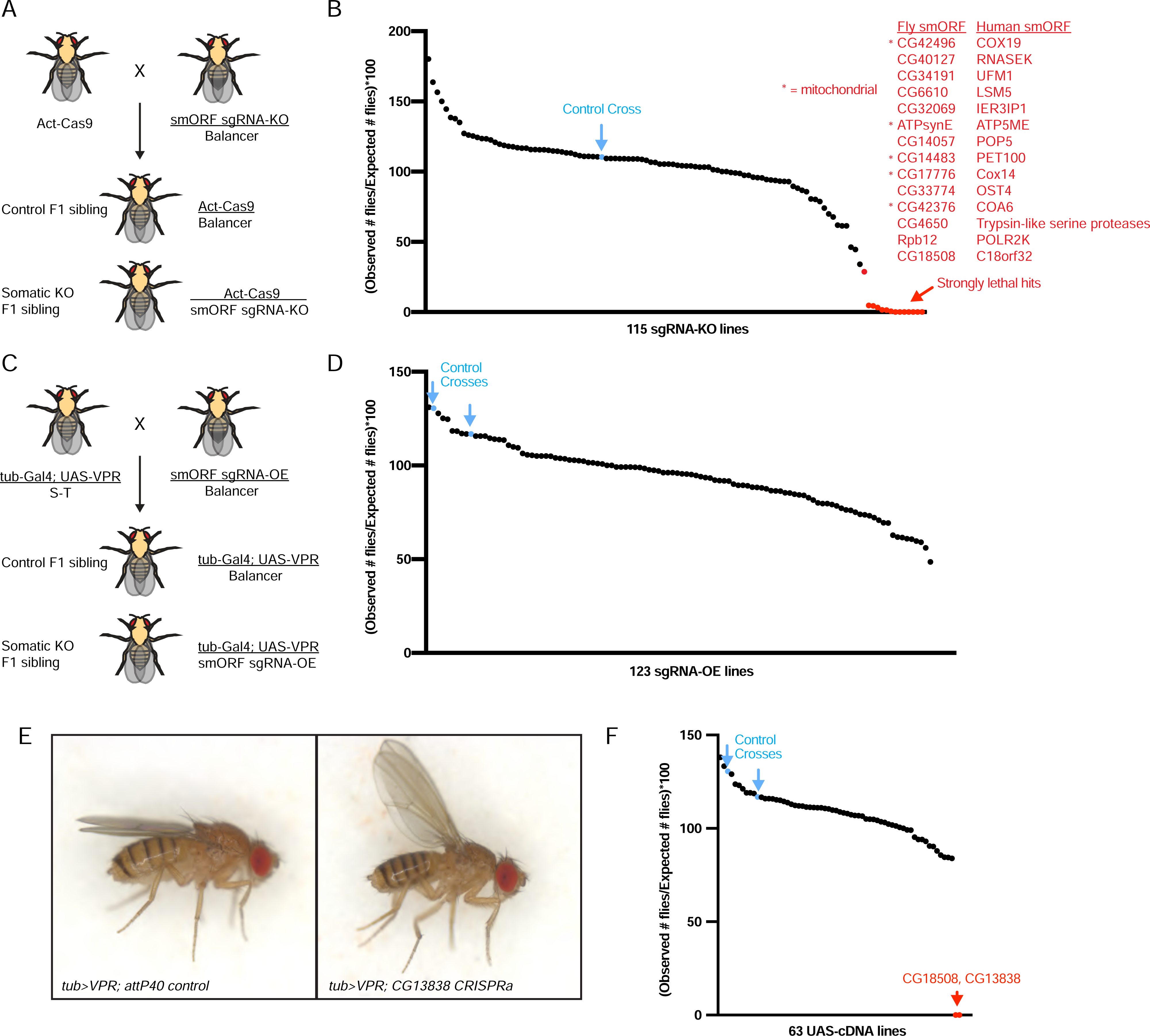
Functional characterization of conserved smORFs by F1 CRISPR in vivo screening. **(*A*)** Genetic cross to perform CRISPR somatic knockout in F1 generation. **(*B*)** Quantification of viability of F1 flies from 115 sgRNA-KO crosses. Number of F1 progeny counted per cross was 918>n>33. **(*C*)** Genetic cross to perform CRISPR gene overexpression in F1 generation. ***(D)*** Quantification of viability of F1 flies from 123 sgRNA-OE crosses. Number of F1 progeny counted per cross was 220>n>56. **(E)** Images of adult female flies aged seven days after eclosion for two indicated genotypes. **(F)** Quantification of viability of F1 flies from 68 UAS-cDNA crosses. Number of F1 progeny countered per cross was 706>n>101.

To determine if the 14 putative essential smORFs play an important role in individual tissues, we crossed the 14 sgRNA-KO hit lines to cell-type specific Cas9 lines (muscles, gut enterocytes, dorsal thorax, wing disc, neurons) (Figure S4). Nearly all sgRNA-KOs reduced viability when expressing Cas9 in neurons (12/14), whereas none reduced viability with muscle Cas9 (0/14). Interestingly, a subset of sgRNA-KO lines had reduced viability only when Cas9 was expressed in neurons (CG14057, CG40127, CG14812, CG17776). In contrast, Rbp12 sgRNA-KO was lethal or showed low viability with all Cas9 lines except muscle, and CG4650 sgRNA-KO did not cause any obvious phenotypes with any of the six tissue-specific Cas9 lines. Finally, using a larval wing disc Cas9 line, seven sgRNA-KO lines caused adult wing defects, such as notching or crumpled wings (Figure S4).

To over-express smORF genes in a systematic manner, we used CRISPR activation (CRISPRa), where a sgRNA targets the promoter region and increases expression of the endogenous gene (sgRNA-OE) via a catalytically dead Cas9 (dCas9) fused with a transcriptional activator (e.g. VPR or SAM) (Ewen-Campen et al., 2017; Jia et al., 2018). Similar to our sgRNA-KO collection, we generated a collection of 197 transgenic sgRNA-OE lines that target 176 smORF genes. Each smORF sgRNA-OE line was crossed with *tub-Gal4*, *UAS-dCas9-VPR* (abbreviated *tub>VPR*), and the F1 progeny were screened for phenotypes as described for the sgRNA-KO collection (Figure 5C; Supplemental File 7). Of the 123 sgRNA-OE lines tested, a small number had reduced numbers of expected progeny, however none were statistically significant when compared to control crosses (Figure 5D). Interestingly, CG13838 sgRNA-OE resulted in viable adults that were flightless and had a “held-up” wing phenotype (Figure 5E). Both phenotypes were 100% penetrant (n=100 flies). None of the remaining tested lines produced aberrant morphological or behavioral phenotypes. To validate overexpression by CRISPRa, we crossed 14 smORF-OE lines to *tub>VPR* and analyzed target gene expression in adults using quantitative PCR (qPCR) (Figure S5). These results showed that 6/14 sgRNA-OE lines had significantly elevated transcript expression when normalized to *Rp49* or *Gapdh* and compared to negative control crosses (*attP40*).

To complement our results with CRISPRa, we overexpressed a subset of smORFs using *UAS-cDNA* lines (Figure 5F; Supplemental File 7), which generally results in higher levels of overexpression (Ewen-Campen et al., 2017). Of the 63 lines tested, representing 58 genes, two lines were lethal, *UAS-CG18508* and *UAS-CG13838*. All other lines had no obvious defects in viability, morphology, or behavior.

### Functional analysis of 25 uncharacterized smORF genes by whole animal KO

F1 CRISPR screening tools are fast and scalable but have technical limitations, namely that CRISPR-KO can produce mosaic phenotypes (Zirin et al., 2020) or phenotypes outside the target tissue (Port and Bullock, 2016). Therefore, we wanted to apply a more robust genetic tool to modify smORF gene function – whole animal knockout. Since this requires greater time and resources, we targeted a subset of the conserved smORFs.

Using a combination of gene function prediction and manual searching (see Materials and Methods), we identified 25 conserved smORF genes with minimal to no previous experimental characterization in any organism with a corresponding homolog (Supplemental File 7). Interestingly, 12 of these smORFs have a paralog in *Drosophila* (Figure S6A). Using CRISPR/Cas9, we generated whole animal KOs for each of the 25 smORF genes and multi-gene KOs for each paralog group (Supplemental File 7; Figure S6B-D). When possible, we generated at least two independently derived alleles for each smORF. The resulting homozygous animals were assessed for mutant phenotypes. Remarkably, nearly all smORF KO lines were viable, fertile, and had no obvious morphological or behavioral defects (Supplemental File 7). Interestingly, KO of the paralogs *CG32736* and *CG42308* was lethal, either as single or double KO (Supplemental File 7; Figure S6B-C). Characterization of *CG32736* and *CG42308* is described elsewhere (Bosch et al., 2022).

We reasoned that viable smORF KO mutants might reveal a mutant phenotype if raised under stressful conditions. To test this, we transferred 24hr old homozygous smORF KO embryos onto modified foods known to cause animal metabolic stress, starvation (Zirin et al., 2015), high fat (Birse et al., 2010), and high salt (Stergiopoulos et al., 2009), and measured developmental timing. On normal food, all tested KO mutants had similar developmental timing compared to wild-type (Figure 6A). However, several KO mutants exhibited significant developmental delays, low viability, or lethality on stressful foods (Figure 6B-D). For example, two independent CG17931-KO alleles were lethal on starvation food and were developmentally delayed or low viability on high salt food. In addition, two independent alleles of CG42371-KO showed low viability on high salt food, and one CG42371 allele had low viability on high fat food. Finally, bc10-KO was developmentally delayed on high salt food.

**Figure 6.**
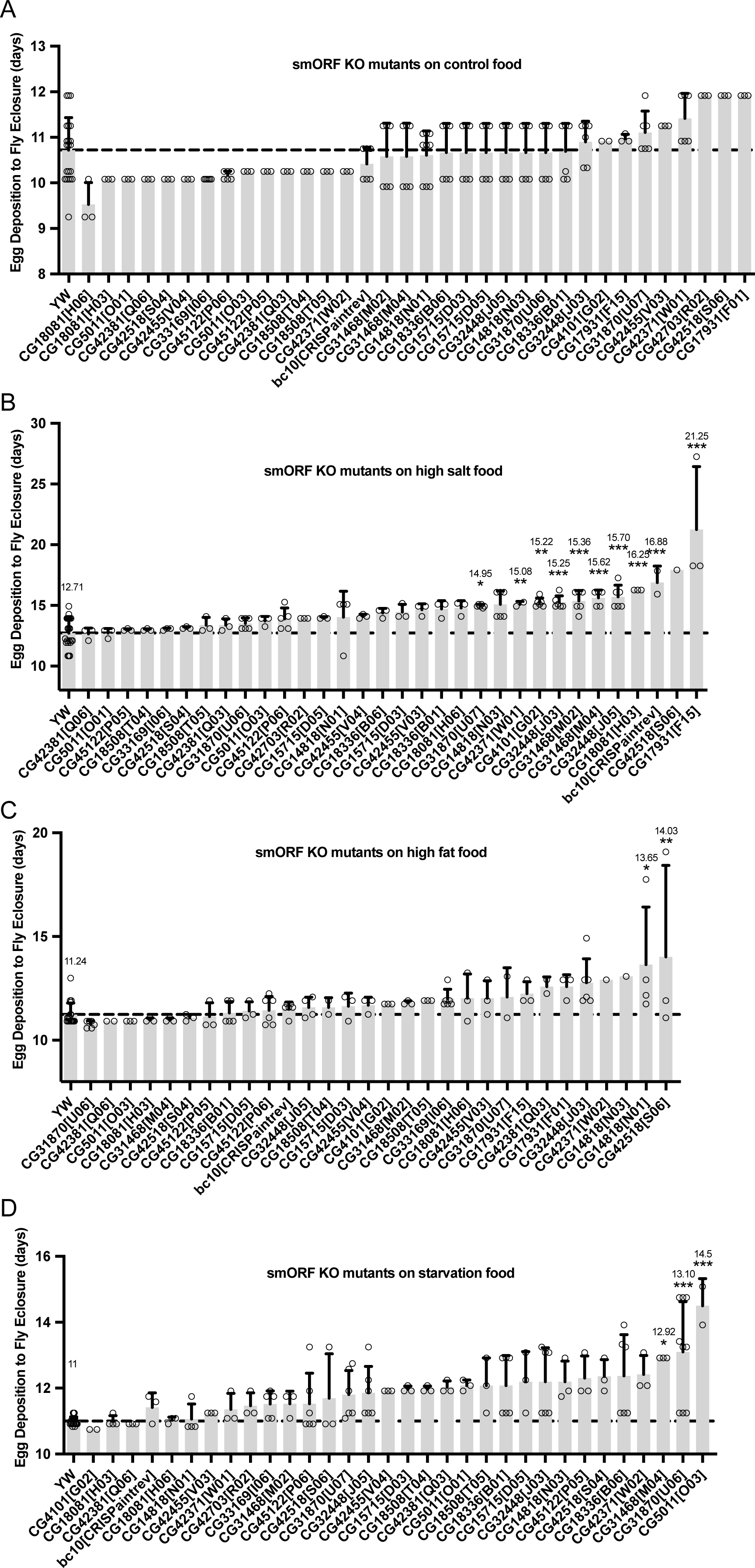
Full CRISPR knockout of 25 uncharacterized smORFs raised on stressful foods. Developmental timing of egg deposition to adult eclosure in smORF KO mutants raised on ***(A)*** control food, ***(B)*** high salt food (30% NaCl), ***(C)*** high fat food (30% coconut oil), ***(D)*** starvation food (30% food, 70% PBS+1%agar). Significance was determined by One Way Anova test followed by a Dunnet Post-Hoc, P=0.05(*), P=0.01(**), P=0.001(***), error bars calculated with SD.

## Discussion

We identified 298 conserved smORFs between humans and fruit flies, the vast majority of which are conserved broadly across bilaterians, with 68 conserved in *Arabidopsis*. The decreased number of smORFs in C.elegans, zebrafish, and Arabidopsis may be due to gene loss or extreme divergence. Ribosome profiling and prior proteomic experiments support the translation of 208 of these smORFs. The remaining 90 without direct evidence of translation are largely highly tissue specific (50 are specific to male reproductive tissues), and likely more targeted samples are needed for detection. Of course, the thresholds we selected for defining smORFs are somewhat arbitrary, and more lenient parameterizations (>100aa) would yield more expansive lists. Certainly, this threshold produced an under-studied collection of genes with diverse functions and phenotypes.

Of the 298 conserved smORFs, 32 are arranged in polycistronic transcripts. However, this gene architecture is rarely conserved in distant species, with only four remaining polycistronic in distant species – the remainder are partitioned into distinct transcriptional units, often on different chromosomes. Polycistronic gene architectures may have been selected against in eukaryotes – possibly as part of antiviral defense, as mRNA viruses often make use of internal ribosome entry sites.

The largest class of conserved smORFs is related to mitochondrial structure and function – including components of the oxidative phosphorylation pathway. Two of these predicted mitochondrial smORFs were recently functionally characterized (Bosch et al., 2022), though it possible that not all function in mitochondria. For example, 22 out of 66 predicted mitochondrial smORF genes are uncharacterized “CGs”.

Similarly, human mitochondria are enriched in smORF peptides (Zhang et al., 2020). Intriguingly, *Drosophila* mitochondrial smORFs exhibit highly tissue specific expression patterns after initial ubiquitous maternal deposition. While mitochondrial functional diversity has been explored in the nervous system (Fecher et al., 2019), this study indicates a far broader diversity in mitochondrial architecture throughout the developing organism. The profound conservation of these genes, along with their tissue-specific expression patterns, indicates that mitochondria are compositionally, and therefore functionally, optimized in a tissue-specific fashion. This observation points to an evolutionary impetus for the translocation of mitochondrial genes to the host nuclear genome -- tissue-specific regulation by host nuclear factors – an intriguing direction for future study.

Results from our F1 CRISPR knockout screens revealed a number of essential smORF genes. Interestingly, animal homologs of these gene hits (10/14) show lethality in other organisms (Marrvel.org) (Wang et al., 2017), and 5/14 are predicted mitochondrial (Supplemental File 2), suggesting that fly lethality may be due to disrupted mitochondrial function. Furthermore, these essential smORFs may be required in different tissues. For example, Rbp12 knockout in either the gut, dorsal thorax, wing disc, or neurons caused significant reduction in animal viability. In contrast, four essential smORFs were only lethal when knocked out in neurons.

One interesting hit from our F1 knockout screen was *CG18508,* the fly homolog of *C18orf32.* The encoded protein has been shown to associate with lipid droplets (Bersuker et al., 2018). While ubiquitous *CG18508* knockout reduced viability, knockout in six tissues did not. However, *CG18508* knockout in the wing disc caused adult wing notching. Surprisingly, homozygous *CG18508* mutants are viable, fertile, and had normal wing morphology, suggesting that *CG18508* sgRNA-KO has off-target effects.

Interestingly, overexpression of *CG18508* by UAS-cDNA was lethal. While it is not clear if *CG18508* overexpression if physiologically relevant, it would be intriguing to examine if animals die due to defects in lipid storage.

Results from the smORF overexpression screen revealed one additional smORF with notable phenotypes. Overexpression of *CG13838* by CRISPRa resulted in flightless adult flies with “held-up” wings, whereas overexpression by UAS-cDNA was lethal. Since UAS-cDNA generally results in higher transcript expression compared to CRISPRa (Ewen-Campen et al., 2017), *UAS-CG13838* cDNA lethality may be due to higher expression. The *C. elegans* homolog of *CG13838*, *bubblin* (*bbln*), has recently been characterized as essential for intermediate filament function (Remmelzwaal et al., 2021). Interestingly, other fly mutations have been described that cause a wing phenotype, such as *heldup*, which disrupts muscle thin filaments (Beall and Fyrberg, 1991). Like intermediate filaments, thin filaments are actin-based cytoskeletal structures (Henderson et al., 2017). Therefore, *CG13838* overexpression may interfere with thin filaments and/or intermediate filaments in muscle.

Many conserved smORFs have known or predicted functions. For example, smORF CG14483 is uncharacterized in *Drosophila*, but its human homolog PET100 (65% aa similarity, 37% aa identity) is a known regulator of mitochondrial complex IV biogenesis and mutated in families with mitochondrial complex IV deficiency nuclear type 12 (MC4DN12) (Lim et al., 2014). In contrast, we identified 25 smORFs with little to no characterization in any organism. For example, CG32736 is homologous to human Small Integral Membrane Protein 4 (SMIM4) (66% aa similarity, 46% aa identity), but had not been experimentally studied in any organism until recently (Bosch et al., 2022; Dennerlein et al., 2021; Liang et al., 2022). Studying poorly characterized smORF genes like *CG32736* could reveal new biology and/or help understand human disease progression. Interestingly, we found three previously uncharacterized smORFs (*CG17931/ SERFs*, *CG42371/ CEBPZOS*, *bc10/BLCAP*) that were required for normal developmental progression on stressful food diets.

## Acknowledgements

We thank Benjamin W Booth for processing the RNA-seq data and submitting RNA and ribosome profiling data to the SRA. We thank Marcus Stoiber for his initial identification of conserved smORFs. We thank Nick Ingolia and Jonathan Weissman for advice about ribosome profiling. This work was funded by an award from the NIH National Human Genome Research Institute (R01HG009352) to S.E.C. (Principal Investigator). J.A.B. was supported by the Damon Runyon Foundation (DRG-2258-16) and a “Training Grant in Genetics” T32 Ruth Kirschstein-National Research Service Award institutional research training grant funded through the NIH/National Institute of General Medical Sciences (T32GM007748). NP is an HHMI investigator. This article is subject to HHMI’s Open Access to Publications policy. HHMI lab heads have previously granted a nonexclusive CC BY 4.0 license to the public and a sublicensable license to HHMI in their research articles. Pursuant to those licenses, the author-accepted manuscript of this article can be made freely available under a CC BY 4.0 license immediately upon publication.

## Materials and Methods

### Bioinformatic identification of 298 fly-human conserved smORFs

First, 266,066 human smORFs were selected, including all annotated human smORF transcripts (18,494 annotated in GENCODE version 24 less than 100aa), and 215,901 smORFs identified by 3-frame translations of all human transcripts that lack long ORFs (11-100aa). This set was filtered to identify high-confidence elements by leveraging a stringent, high-confidence set of conserved *Drosophila* smORFs. In *Drosophila*, there are 960 genes encoding unique peptides with no more than 100 aa from protein-coding genes annotated at FlyBase (release 6.49) with evidence of translation. This set was expanded by taking smORFs (11-100aa) predicted from two independent studies (Aspden et al., 2014), adding 2,819 additional smORFs with evidence of translation from either Ribosome Profiling or conservation among Drosophilidae.

For our ortholog discovery workflow, see Fig. 1A. Specifically, we used Diopt v8 and 9 (https://www.flyrnai.org/cgi-bin/DRSC_orthologs.pl) for ortholog analysis. In parallel and to corroborate results, we also used deltablast 2.9.0+ build Sep 30 2019 01:57:31 with the following parameters: (Matrix : BLOSUM62); (Gap Penalties: Existence: 11, Extension :1); (Neighboring words threshold: 11); (Window for multiple hits:40). We then filtered the deltablast results for *D. rerio*, *C. elegans*, *A. thaliana* proteins using the following cutoffs <=250aa and E-value <=10^-1^. The *D. rerio*, *C. elegans* and *A. thaliana* peptide sequence files used, are as follow: Danio_rerio.GRCz11.pep.all.fa, <u>Caenorhabditis_elegans.WBcel235.pep.all.fa</u> (https://ftp.ensemblgenomes.ebi.ac.uk/pub/metazoa/release-56/fasta/caenorhabditis_elegans/) and TAIR10_pep_20101214.faa (https://www.arabidopsis.org/download/index-auto.jsp?dir=%2Fdownload_files%2FProteins%2FTAIR10_protein_lists). These analyses identified orthologs in human for 291 fly genes, and an additional 7 were discovered using sim3 analysis (Chao et al., 1997).

### Clustering of ontological anatomical annotations of embryonic expression patterns

Embryos were clustered using a bagged and cross validated procedure to ensure cluster stability. The importance of stability was made apparent to us when, during the course of our study, one additional smORF was discovered and added to this analysis, and the resulting clusters differed significantly. We stabilized the clustering as follows:

First, we subsampled 80% of smORFs and used hierarchical clustering with Ward linkage to form candidate clusters, holding out 20%. Here, the hold-out is to assess stability by inducing some randomness -- note that this differs from the use of a holdout in supervised learning. We selected 10K 80/20 splits. Cluster number was selected using the Gap Statistic -- the first local maximum value was selected. We computed a proximity matrix for consensus clustering as follows: for each pair of smORFs we recorded the fraction of our 10K bagged clusterings in which they appeared in the same cluster. This matrix was then input to hierarchical clustering with Ward linkage, and the Gap Statistic was used to select cluster number as above.

### Embryo Collections for Coordinated Ribosome and RNA-Seq Profiling

Embryos from ∼14 g of Oregon-R flies were collected on standard molasses collection trays after flies were acclimated to the environmental conditions of the cage (27℃ and 70% humidity) for three days. Six, two-hour embryonic time periods (0-2, 2–4 hr., 4–6 hr., 10-12 hr., 14–16 hr., and 16–18 hr. were collected simultaneously, which allowed for immediate RNA-seq library and ribosome profiling construction of all six stages. See Supplemental File 6 for detailed the ribosome profiling library protocol.

### RNA Preparation and Sequencing Methods

Embryos were homogenized using a Pellet Pestle Cordless Motor (Kimble Cat. No. 749540-0000; Pellet Pestles Sigma Cat. No. Z359947), RNA was extracted using TRIzol Reagent (Thermo Fisher, Cat. No. 15596026) and purified with the RNeasy Mini Kit (QIAGEN Cat. No. 74106). Libraries were constructed with the NEBNext Ultra Directional RNA Library Prep Kit for Illumina according to manufacturer’s recommendations (NEB, cat. no. E7420) using 14 cycles of PCR. Libraries were sequenced on the Illumina NovaSeq 6000. Raw sequencing data is available at the NCBI Short Read Archive (SRA): SRR18575339, SRR18575340, SRR18575342, SRR18575343, and SRR18575345 and SRR18575346.

We used the STAR aligner v2.73a to align RNA-seq data to the *D. melanogaster* genome (Rel 6). The picard 2.20.1 MarkDuplicates tool was used to remove PCR duplicates and the deduplicated BAM alignment files were converted to bigWig format using a custom tool and the UCSC bedGraphToBigWig tool. We ran fastp 0.20.1 to get FASTQ file statistics.

### Ribosome Profiling Methods

Polysome profiling was performed on all six samples. Briefly, embryos from each time period (i.e., sample) were treated with harringtonine (LKT Laboratories, H0169) in mild lysis buffer followed by the addition of cycloheximide and immediate grinding. Samples were subjected to a 10%-50% sucrose gradient and all polysomes were collected and combined. Collected polysome fractions were pelleted via sucrose cushion (34%) and subjected to RNaseI digestion. After resuspension another cushion (34%) was performed and RNA was coprecipitated with GlycoBlue (Thermo Fisher, cat. No. AM95150). Recovered RNA was then run on a 15% TBE-Urea Gel (Thermo Fisher cat. no. EC68852BOX), followed by gel size selection to isolate ribosome protected fragments (26-31 nt) (Zymo Research, cat. no. R1070). Following end-repair phosphorylation of RNA molecules, libraries were constructed with the NEBNext Small RNA Library Preparation Kit according to manufacturer’s recommendations (NEB, cat. no. E7330). See Supplemental File 6 for detailed protocol.

### Sequencing, read processing and mapping

Ribo-seq libraries were sequenced with the Illumina NovaSeq 6000. Raw sequencing data is available at the SRA: SRR18575338, SRR18575341, SRR18575344, SRR18575347, SRR18575348, and SRR18575353. Read processing and mapping were performed on an Ubuntu 18.04 Linux cluster running Kubernetes v1.16. on 240 total cores with 1500 GB of total RAM. Reads were processed with the following commands:

I QC of raw reads with FastQC (http://www.bioinformatics.babraham.ac.uk/projects/fastqc/) fastqc file_one.fastq.gz file_two.fastq.gz
II. Clip adapters from Illumina raw reads with the following commands with Cutadapt (Martin, 2011) cutadapt -a adapter sequence for file_one.fastq.gz \ -A adapter sequence for file_one.fastq.gz -j 10 \ -o file_one_clipped.fastq -p file_two_clipped.fastq \ file_one.fastq.gz file_two.fastq.gz
III. Trim clipped reads based on position quality with the following commands: cutadapt -q 33 -j 12 -o file_one_clipped_trimmed.fastq file_one_clipped.fastq cutadapt -q 33 -j 12 -o file_two_clipped_trimmed.fastq file_two_clipped.fastq
IV. QC of processed reads: fastqc file_one_clipped_trimmed.fastq \ file_two_clipped_trimmed.fastq
V. Remove reads smaller than 26 bp and larger than 31 bp: cutadapt --pair-filter=any --minimum-length=26 --maximum-length=31 -j 20 \ -o file_one_clipped_trimmed_min_max_removed.fastq \ -p file_two_clipped_trimmed_min_max_removed.fastq \ file_one_clipped_trimmed.fastq file_two_clipped_trimmed.fastq
VI. Map first to the rDNA reference with Bowtie2 (Langmead and Salzberg, 2012) to 1) remove rRNA sequences and decrease mapping time to nuclear reference genome, and 2) assess the percentage of rRNA contamination in each library: Bowtie2 -x rDNA_reference_directory \ -1 file_one_clipped_trimmed_min_max_removed.fastq \ -2 file_two_clipped_trimmed_min_max_removed.fastq \ --seedlen 12 --un-conc Bowtie2_mapping_directory -p 12 \ -S rDNA_mapping.sam
VII. Map non-rRNA reads to nuclear reference genome with STAR (Dobin et al., 2013): STAR --runThreadN 25 --genomeDir --outFileNamePrefix \ --outSAMtype BAM SortedByCoordinate \ --winAnchorMultimapNmax 100 --seedSearchStartLmax 20 \ --outFilterMismatchNmax 3 --readFilesIn \ Bowtie2_mapping_directory/un-conc-mate.1 \ Bowtie2_mapping_directory/un-conc-mate.2

### Detection of smORFs with Ribosome Profiling

Identification of translated sequences was performed using ORFquant (Calviello et al., 2020). For each predicted ORF with mapped reads we recorded all ORFquant summary statistics. We classified ORFs as detected if the adjusted P-value was less than or equal to 0.05 (Supplemental Table 5).

### Gene enrichment analyses

GO and KEGG enrichment analyses were performed with g:Profiler (Raudvere et al., 2019).

### Molecular biology

Fly genomic DNA was isolated by grinding a single fly in 50µl squishing buffer (10 mM Tris-Cl pH 8.2, 1 mM EDTA, 25 mM NaCl) with 200µg/ml Proteinase K (3115879001, Roche), incubating at 37°C for 30 min, and 95°C for 2 minutes. PCR was performed using Taq polymerase (TAKR001C, ClonTech) when running DNA fragments on a gel, and Phusion polymerase (M-0530, NEB) was used when DNA fragments were sequenced or used for molecular cloning. DNA fragments were run on a 1% agarose gel for imaging or purified on QIAquick columns (28115, Qiagen) for sequencing analysis. Sanger sequencing was performed at the DF/HCC DNA Resource Core facility and chromatograms were analyzed using Lasergene 13 software (DNASTAR).

For isolating flies with frameshift indels in smORF genes, the target site was PCR amplified from single fly genomic DNA and PCR fragments were Sanger sequenced. For isolating flies with whole gene deletion of smORF genes by dual sgRNA cutting, the target region was PCR amplified from single fly genomic DNA. Primers were designed to flank the two sgRNA cut sites, such that a deletion of the intervening sequence would produce a clear band size difference on an agarose gel. Deletion PCR fragments were Sanger sequenced. Genotyping primer sequences are listed in Supplemental File 7.

For RT-qPCR analysis of smORF overexpression by CRISPRa, adult flies (tub-Gal4, UAS-dCas9-VPR, sgRNA-OE) were flash frozen in liquid nitrogen. 4-14 frozen flies (equal mixture of males and females) were homogenized in 1000ul Trizol (Invitrogen 15596026), RNA partially purified by chloroform extraction, and RNA extracted using a Direct-zol RNA Miniprep kit (Zymo Research, R2050). cDNA was generated using the iScript Reverse Transcription Supermix (BioRad 1708840). cDNA was analyzed by RT-qPCR using iQ SYBR Green Supermix (BioRad 170-8880) on a CFX96 Real-Time system (BioRad). qPCR primer sequences are listed in Supplemental File 7. Each qPCR reaction was performed with five biological replicates (except *CG14818*, which had two biological replicates), with two technical replicates each. Data from smORF specific primers were normalized to primers that amplify GAPDH. Statistical significance was calculated using a T-Test.

### Molecular cloning

Plasmid DNAs were constructed and propagated using standard protocols. Briefly, chemically competent TOP10 E.coli. (Invitrogen, C404010) were transformed with plasmids containing either Ampicillin or Kanamycin resistance genes and were selected on LB-Agar plates with 100µg/ml Ampicillin or 50µg/ml Kanamycin. Oligo sequences are in Supplemental File 8.

#### sgRNA expression plasmids

Plasmids encoding sgRNAs were generated using previously described protocols. sgRNAs were designed using the Find CRISPR tool (https://www.flyrnai.org/crispr3/web) for optimal predicted cutting activity (Zirin et al., 2020). For sgRNAs cloned into *pCFD3* (Port et al., 2014), annealed oligos encoding a sgRNA spacer were ligated into *pCFD3* digested with BbsI (NEB, R3539). For sgRNAs cloned into *pCFD4* (Port et al., 2014), dual sgRNAs were PCR amplified from *pCFD4* template, and inserted by Gibson assembly (NEB, E2611) into *pCFD4* digested with BbsI. For sgRNAs cloned into *pCFD5* (Port and Bullock, 2016), dual sgRNAs were PCR amplified from *pCFD5* template, and inserted by Gibson assembly into *pCFD5* digested with BbsI. sgRNAs cloned into *pCFD3* and were performed by DRSC/TRiP (https://fgr.hms.harvard.edu/). Information on sgRNA-KO and sgRNA-OE plasmids generated by the DRSC/TRiP is available at https://www.flyrnai.org/tools/grna_tracker/web/. Information on remaining *pCFD4* and *pCFD5* sgRNA plasmids is in Supplemental File 8.

#### pEntr plasmids

Entry plasmids were generated by PCR amplifying coding sequence and inserting into *pEntr* using either dTopo (Invitrogen, K240020) or Gibson assembly (NEB, E2611). cDNA sequence was PCR amplified from cDNA reverse transcribed from total RNA (either S2R+ cell or adult fly) or adult fly genomic DNA.

#### Gateway cloning LR reactions

Gateway cloning reactions were performed using LR Clonase II Enzyme mix (Invitrogen 11791-020). pEntr cDNA plasmids were recombined with *pWalium10-roe* (Perkins et al., 2015).

### Fly Genetics

Flies were maintained on standard fly food at 25°C unless otherwise noted.

Fly stocks were obtained from the Perrimon lab collection, Bloomington Stock center (indicated with BL#), or generated in this study (see below).

Perrimon Lab stocks:

*yv; Gla/CyO*

*yw; nos-Cas9attP40/CyO*

*yw;; nos-Cas9attP2*

*lethal/FM7,GFP*

*yw; Gla/CyO*

*yw;; TM3, Sb/TM6b*

*lethal/FM7,GFP;; TM3, Ser*

*yw; Sp/CyO; MKRS/TM6B*

Bloomington Stocks:

sgRNA lines (see Supplemental File 8)

*UAS-dCas9-VPR; tub-Gal4/S-T* (*tub>VPR*) (BL67048)

*Actin-Cas9* (BL54590)

*attP40* (BL36304)

*Mhc>Cas9* (BL67079)

*LSP>Cas9* (BL67087)

*Myo1a>Cas9* (BL67088)

*Pnr>Cas9* (BL67077)

*Nub>Cas9* (BL67086)

*Elav>Cas9* (BL67073)

*yw* (BL1495)

Information on the smORF KO and UAS-cDNA stocks are described in Supplemental File 8, including Bloomington Stock #s.

#### smORF knockout by frameshift indel by transgenic crossing

Flies expressing a sgRNA that targets the 5’ coding sequence (see Supplemental File 8) were crossed with *nos-Cas9* flies. *nos-Cas9attP2* was used for targeting genes on Chromosomes X and II, and *nos-Cas9attP40* was used for targeting genes on chromosome III. F1 progeny were crossed with a balancer strain. Single fly F2 progeny were crossed with a balancer strain, taken for genotyping, and F2 crosses with a frameshift indel were kept and balanced and homozygosed if possible. Frameshift knockout lines were generated either by WellGenetics, Shu Kondo, or in the Perrimon lab.

#### smORF knockout by full gene deletion by injection

Plasmids encoding two sgRNAs that flank a gene locus were injected into *nos-Cas9* embryos. Injected F0 adults were crossed to with a balancer strain. Single fly F1 progeny were crossed with a balancer strain, taken for genotyping, and F1 crosses with a full gene deletion were kept and balanced and homozygosed if possible.

#### smORF knockout by CRISPaint insertion by injection

To generate the bc10 KO allele, a plasmid encoding a sgRNA that targets the 5’ coding sequence of bc10 (GP01409) was co-injected with pCRISPaint-T2A-Gal4-3xP3-RFP (Addgene #127556) into *nos-Cas9attP40* embryos. Injected F0 adults were outcrossed to *yw*, single RFP+ F1 progeny were crossed with *yw;; TM3, Sb/TM6b,* and the insertion in bc10 was verified by PCR and sanger sequencing. The *bc10-CRISPaint* allele contains *T2A-Gal4* inserted in the reverse orientation relative to the 5’-3’ *bc10* transcript, and thus is not a Gal4 reporter allele.

To generate double smORF knockout lines, smORF alleles on different chromosomes were brought into the same strain by outcrossing to double balancer lines.

#### CRISPR-KO F1 crosses and phenotyping

Of the 165 smORF genes targeted with at least sgRNAs for knockout, 11 genes were tested with two independent sgRNAs, and four genes were tested with three sgRNAs. Lines expressing sgRNAs that target smORF 5’ coding sequence were crossed with a line ubiquitously expressing Cas9, *Act5c-Cas9*. Specifically, male sgRNA flies were used that were heterozygous with a balancer chromosome (CyO or TM3, Sb), and were crossed with homozygous *Act5c-Cas9* female flies. To quantify the viability of F1 flies with somatic KO, we recorded the number of balancer progeny (*Act5c-Cas9/Bal*) and non-balancer progeny (*sgRNA/Act5c-Cas9*). The total number of F1 progeny counted per cross was 920>n>31. To calculate a viability score, the number non-balancer flies was divided by the mendelian expected number of non-balancer flies (# F1 progeny/2) and multiplied by 100 (# observed non-balancer/# expected non-balancer*100). Negative control crosses were *attp40/CyO* males crossed with *Act5c-Cas9* females. For crosses leading to reduced viability, a Chi-square test was used to determine significance by comparing the # expected non-balancer flies from negative control (attp40) vs experimental (sgRNA) crosses. Those sgRNA hits that had significant reduced viability (Figure 5, Supplemental File 7) were crossed with tissue specific Cas9 lines (Figure S4). Viability scores and Chi-square tests were performed similarly to *Act5c-Cas9* crosses.

#### CRISPRa F1 crosses and phenotyping

Of the 176 smORF genes targeted with at least sgRNAs for overexpression, 19 genes were tested with two independent sgRNAs, and one gene was tested with three sgRNAs. Lines expressing sgRNAs that target upstream of a smORF transcriptional start site (TSS) were crossed with line ubiquitously expressing dCas9-VPR, *tub>VPR* (*tub-Gal4, UAS-dCas9-VPR/S-T*). “S-T” are a second and third chromosome balancer pair that segregate together due to a reciprocal translocation, and are marked by Cy, Hu, and Tb (T(2;3)TSTL14, SM5: TM6B, Tb[1]). Male sgRNA flies were crossed with tub>VPR females. A viability score was calculated similar to CRISPR-KO F1 crosses, (# observed non-balancer/# expected non-balancer*100). When using homozygous sgRNA males, the expected number of non-balancer flies was # F1 progeny/2. When using heterozygous sgRNA/Bal males, the expected number of non-balancer flies was # F1 progeny/4. The total number of F1 progeny counted per cross was 346>n>56. Negative control crosses were *attp40/CyO* males crossed with *tub>VPR* females. Chi-square analysis was performed similarly to CRISPR-KO F1 crosses. No crosses had significantly reduced viability.

#### cDNA overexpression crosses and phenotyping

Transgenic UAS-cDNA lines were crossed with a ubiquitous Gal4 line. For convenience, we used the same driver used for CRISPRa crosses, *tub>VPR* (*tub-Gal4, UAS-dCas9-VPR/S-T*). A viability score was calculated similar to CRISPRa F1 crosses. Negative control crosses were *attp40/CyO* males crossed with *tub>VPR* females. The total number of F1 progeny counted per cross was 706>n>100. Chi-square analysis was performed similarly to CRISPR-KO F1 crosses.

#### Stressful food recipes

For all food types, standard lab fly food was melted in a microwave, and distilled water (dH20) was added as 10% boiled volume (100ml boiled food + 10ml dH20) to replace the evaporated water. This is used as control food. For high salt food, solid NaCl (Fisher Scientific, S271) was added at 30% weight per volume of control food (e.g. 100ml control food + 30g NaCl) and mixed well. For high fat food, solid coconut oil (Sigma, W530155) was added at 30% weight per volume of control food (e.g. 100ml control food + 30g coconut oil) and mixed well. For starvation food, control food was diluted to 30% in 1% melted agar (BD, 214030) in 1x PBS (Gibco, 10010-023) (e.g. 30ml control food + 70ml melted 1% agar in 1x PBS). Melted liquid food types were poured into empty vials and cooled at 4°C.

#### Quantification of developmental timing on stressful food

24hr larvae from homozygous viable smORF lines were transferred to stressful food or control food and raised on this food until pupal eclosion as adults. To increase the fecundity of the adult flies, four days prior to the 24hr larval collection, ∼150 adult flies from each KO line were transferred to fresh food containing yeast paste. Adult flies were transferred onto fresh food bottles containing yeast paste and allowed to lay for 4hr. 24hr after the end of egg deposition, 30 freshly hatched larvae (24hr-28hr old) were transferred into vials containing either control food or stressful food (control, high fat, high salt, starvation). The time of pupariation and fly eclosion was determined once at least 15 flies pupariated and eclosed, respectively. yw flies were included as a negative control genotype. Each genotype-foodtype experiment was carried out in at least triplicate. For those genotype-foodtypes with a developmental delay, significance was calculated using a One-Way ANOVA test run with a Dunnet post-hoc test using GraphPad Prism.

#### Adult wing mounting, imaging, analysis

sgRNA-KO lines were crossed with *nub>Cas9* and the wings of adult progeny were removed using forceps under a dissecting microscope. For each genotype, at least 6 wings were collected. Removed wings were placed onto a drop of mounting medium (50% Permount (Fisher Scientific, SP15), 50% Xylenes (Fisher Scientific X5)) on a microscope slide (Thermo Scientific, 3050) and mounted using a coverslip (VWR, 48393059). The coverslip was sealed to the microscope slide with clear nail polish. Images of the wings were taken using a stereo microscope (Zeiss Axio Zoom V16) at 32x magnification. The area of each wing was measured using the polygon function in ImageJ. The area measurements were analyzed in Graphpad Prism. A One-way ANOVA test was carried out with a Dunnett Post-Hoc test to determine the significance of the area of the wings of the experimental cross progeny when compared with the control cross progeny (*attP40 X nub>Cas9)*.

### Bioinformatics and literature searching

For protein alignments in Figure S6, we downloaded protein sequence files from Flybase.org or NCBI, aligned them using Clustal Omega (https://www.ebi.ac.uk/Tools/msa/clustalo), and generated alignment images using JalView (https://www.jalview.org/).

For literature searching for smORF homolog characterization, we queried every ultraconserved *Drosophila* smORF using the following online tools: Gene2function (Gene2function.org), HUGO Gene Nomenclature Committee (genenames.org), Interpro (ebi.ac.uk/interpro), and Alliance of Genome Resources (alliancegenome.org). Those smORF genes (25) that had no or minimal characterization in orthologs were selected.

