## Supplemental Figures for "Molecular and functional characterization of the *Drosophila melanogaster* conserved smORFome"

A. Complexes of the Respiratory Chain

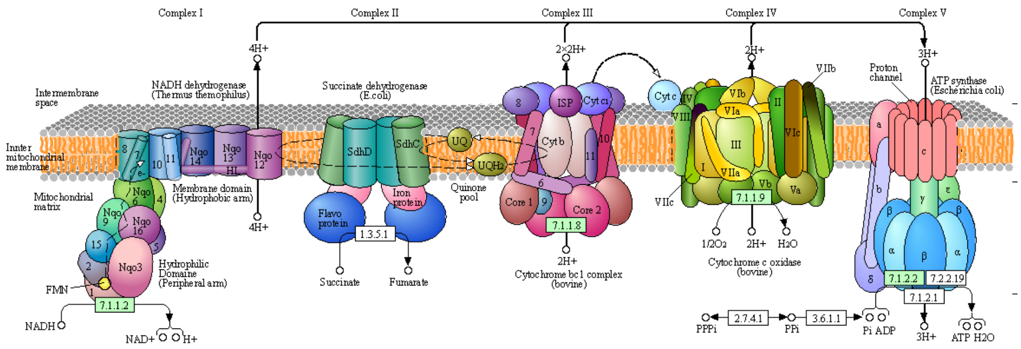

B. *NADH dehydrogenase* (Complex I)

| | | | | | | | | | | | | |
|---|---|---|---|---|---|---|---|---|---|---|---|---|
| ND1 | ND2 | ND3 | ND4 | ND4L | ND5 | ND6 | | | | | | |
| Ndufs1 | Ndufs2 | Ndufs3 | Ndufs4 | Ndufs5 | Ndufs6 | Ndufs7 | Ndufs8 | Ndufv1 | Ndufv2 | Ndufv3 | | |
| Ndufa1 | Ndufa2 | Ndufa3 | Ndufa5 | Ndufa6 | Ndufa7* | Ndufa8 | Ndufa9 | Ndufa10 | Ndufab1 | Ndufa11 | Ndufa12 | Ndufa13 |
| Ndufb1 | Ndufb2* | Ndufb3 | Ndufb4 | Ndufb5 | Ndufb6 | Ndufb7 | Ndufb8 | Ndufb9 | Ndufb10 | Ndufb11 | Ndufc1 | Ndufc2 |

C. Cytochrome c oxidase structural components (Complex IV)

| | | | | | | | | | | | | | |
|---|---|---|---|---|---|---|---|---|---|---|---|---|---|
| COX1 | COX2 | COX3 | COX4I1 | COX5A | COX5B | COX6A | COX6B | COX6C* | COX7A* | COX7B | COX7C* | COX8 | Ndufa4 |

D. F-type ATPase (Complex V)

| | | | | | |
|---|---|---|---|---|---|
| alpha | beta | gamma | delta | epsilon* | |
| OSCP | a | b | c | d | e |
| f | g | f6/h | j | k* | 8 |

E. Cytochrome c reductase (Complex III)

| | | | | | | |
|---|---|---|---|---|---|---|
| UQCRFS1 | CYTB | CYC1 | | | | |
| SYCP3 | UQCR2 | UQCRH* | UQCRB | UQCRQ | UQCR10 | UQCR11* |

**Figure S1.** Conserved smORFs of the Respiratory Chain. A) Electron transport takes place in the mitochondria and is the site of oxidative phosphorylation in eukaryotes. The complexes I to V and the enzymes required for human mitochondrial function are diagramed (Kyoto Encyclopedia of Genes and Genomes (Kanehisa et al., 2022); KEGG). B) Part of Complex I, NADH dehydrogenases are used in the electron transport chain for generation of ATP and to convert NADH to NAD+. Of the forty-nine members all but two (Ndufa3, Ndufc1) have flies orthologs Six are conserved smORFs in flies and humans (green shading). In flies there are two orthologs for the human gene, Ndufb2a (asterisk) orthologs CG40472 (94aa) and ND-AGGG (54aa). C) Complex IV or Cytochrome C Oxidase is a large protein complex. Of the fourteen subunits of the structural components of the CIV complex (Zong et al., 2018), seven are conserved fly and human smORFs (green shading). In flies there are two orthologs for the human genes COX6C, and COX7C. Their orthologs are cype and CG15386 and COX7C and COX7CL, respectively. COX7A is a more complex case with four human variants, COX7A1, COX7A2, COX7A2 and COX7A2P2 with orthology to two smORF fly genes, COX7A and COX7AL2. D) Part of Complex V, ATPases produce ATP from ADP in the presence of a proton gradient. Five are conserved smORFs. The human F-type ATPase epsilon genes (ATP5F1E and ATP5F1EP2) have two fly orthologs ATPsynepsilonL (64aa) and sun (57aa) both fly genes more similar to ATP5F1E than to ATP5F1EP2. The human ATPase k gene has two fly orthologs Neb-cGP and CG15458. E) Part of Complex III, cytochrome C reductases pass electrons to Cytochrome C. Of the ten members four are smORFs, UQCRH, UQCRQ, UQCR10 and UQCR11. Two of the orthologous smORFs, human UQCRH and UQCR11 and are encoded in flies by paralogous genes: UQCR11; UQCR11L and UQCR6.4; UQCR6.4L, respectively. In flies, the second copy, those designated with “L” are uniquely expressed in testes.

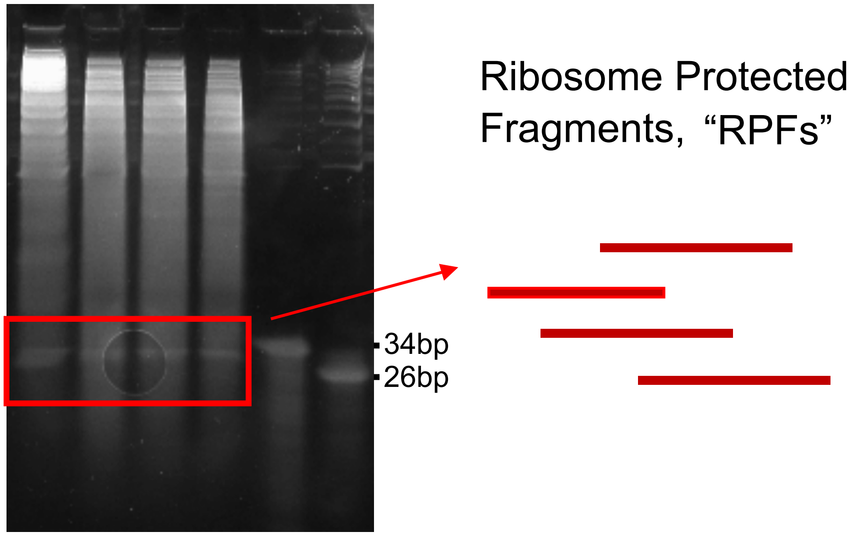

**Figure S2. Size selection on PAGE-Urea Gel (26-34bp):** Recovered RNA was run on a page urea gel and RNA molecules in the range of ribosome protected fragments (RPFs) are excised and purified. After sequencing, the number of in frame reads are analyzed

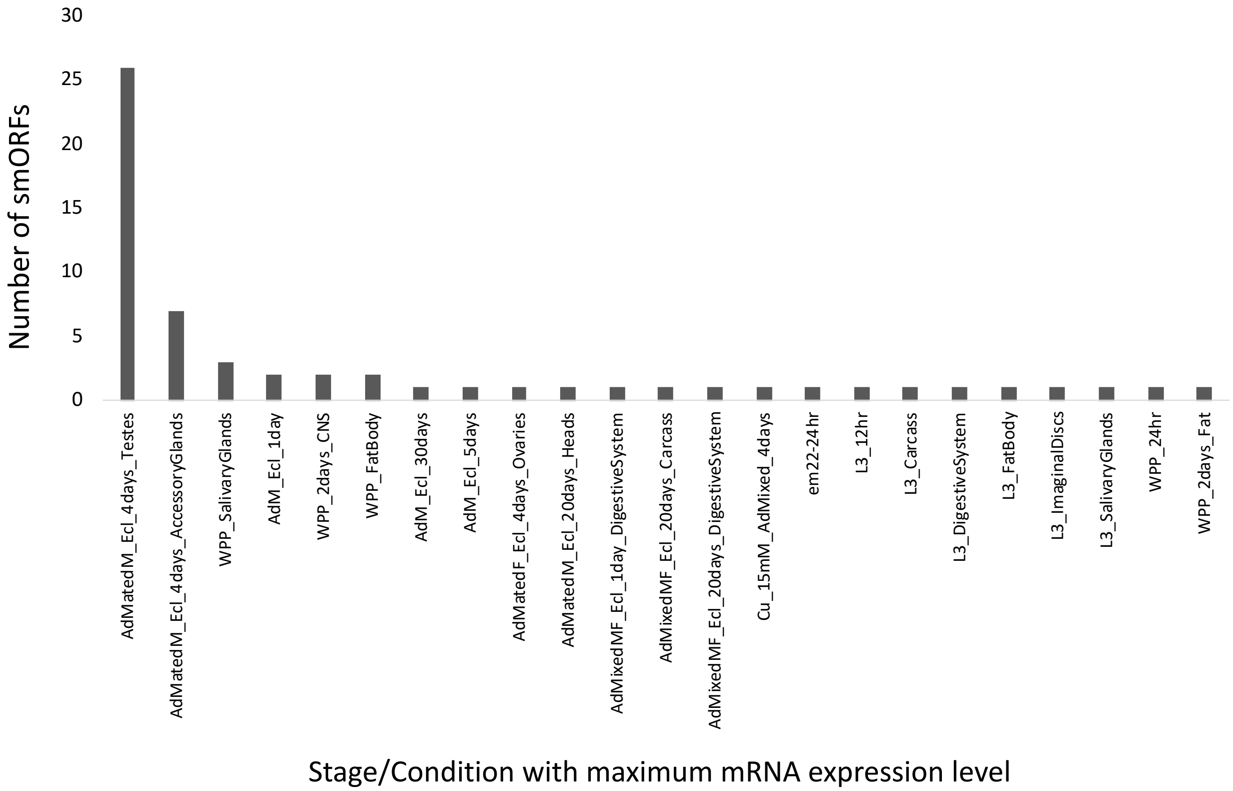

**Figure S3.** The developmental stage/condition of maximum mRNA gene expression (modENCODE) are plotted for ucsmORFs without Ribo-seq or proteomics support.

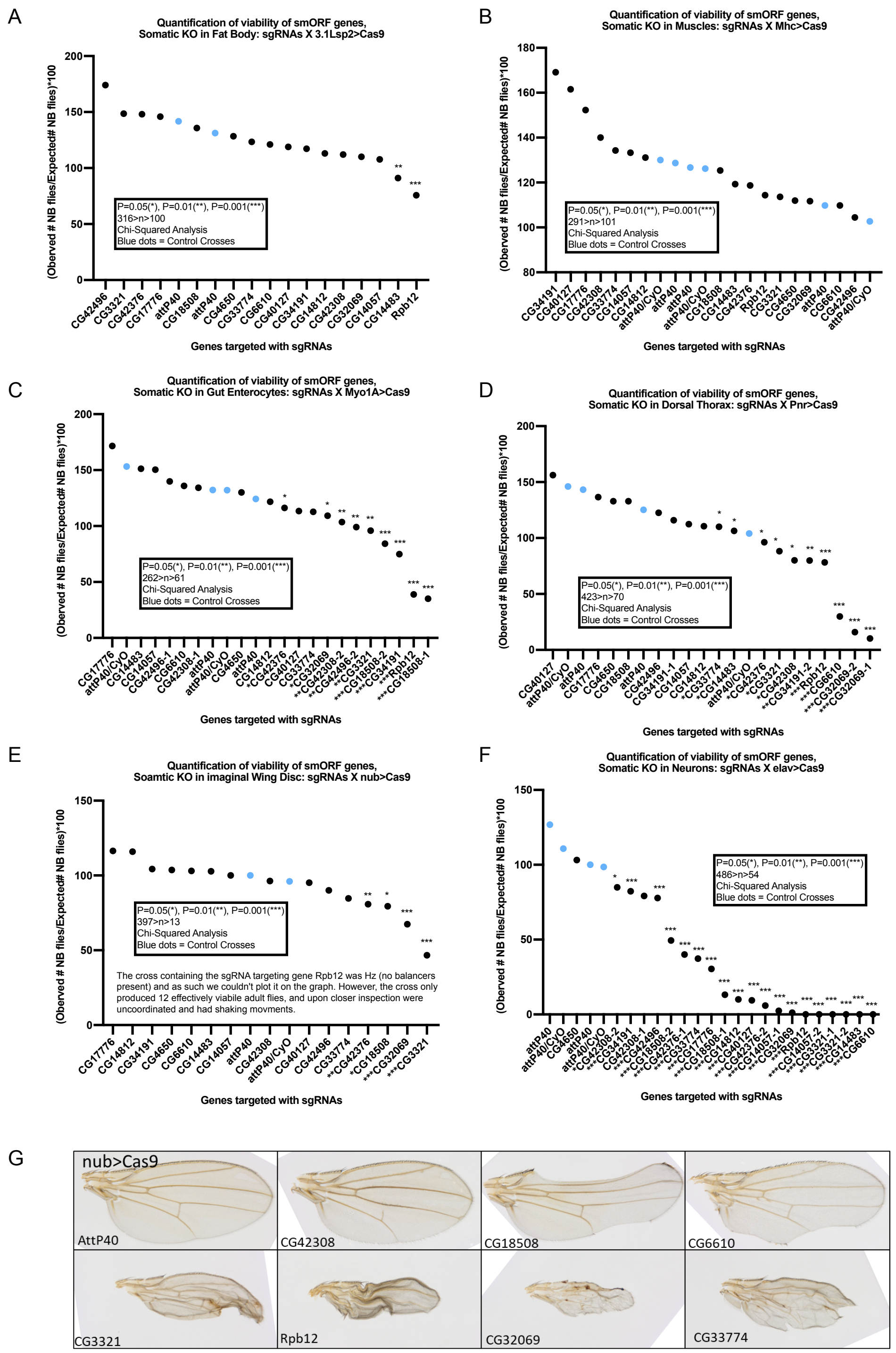

**Figure S4 – Tissue-specific somatic knockout of lethal smORF hits in Figure 1B.** Quantification of viability from CRISPR somatic knock out in **A)** fat body using 3.1Lsp2>Cas9, **B)** muscle using Mhc>Cas9, **C)** gut enterocytes using Myo1A>Cas9, **D)** dorsal thorax using pnr>Cas9, **E)** wing disc using nub>Cas9, or **F)** neurons using elav>Cas9. Significance calculated with a Chi-Squared Test. P=0.05(*), P=0.01(**), P=0.001(***) **G)** Images of adult wings with somatic KO.

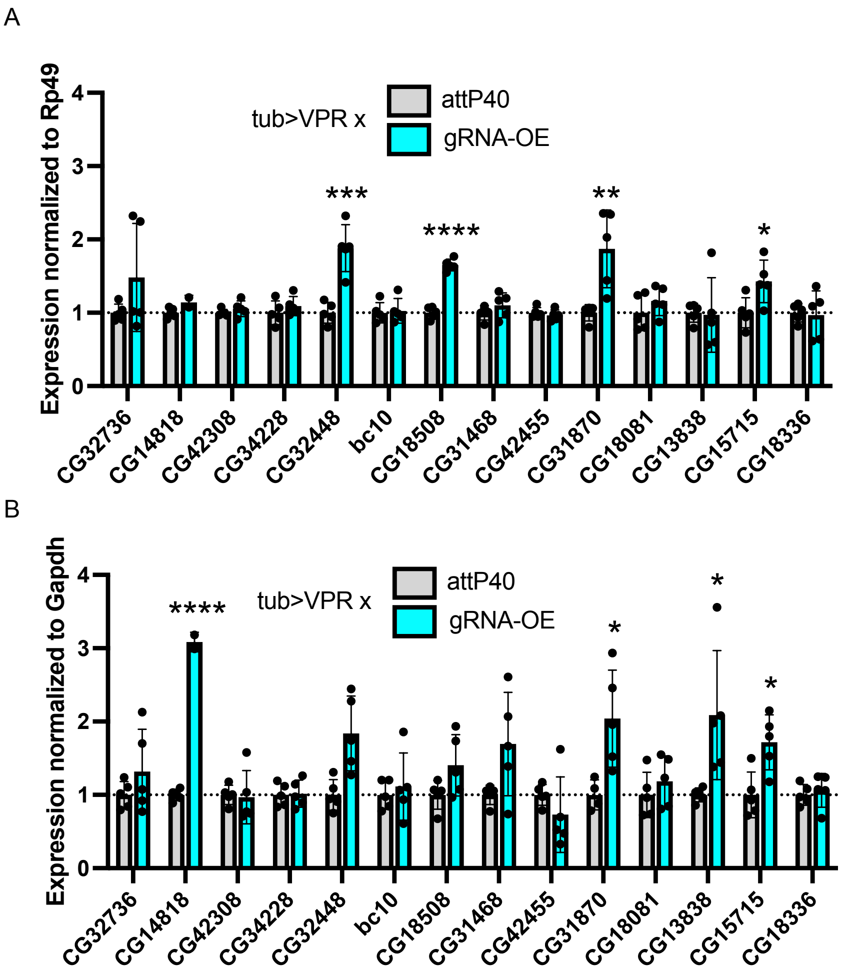

**Figure S5 – qPCR verification of CRISPRa lines.** smORF transcript expression normalized to (A) *Rp49* and (B) *Gapdh* for indicated genotypes. Each sgRNA-OE expression data is normalized to negative control (attP40) expression data. Significance between sgRNA-OE and attP40 control expression levels determined by a T-test. For each genotype, N=5 biological replicates, except CG14818 sgRNA-OE which had 2 biological replicates. P=0.05(*), P=0.01(**), P=0.001(***), P ≤ 0.0001 (****)

A

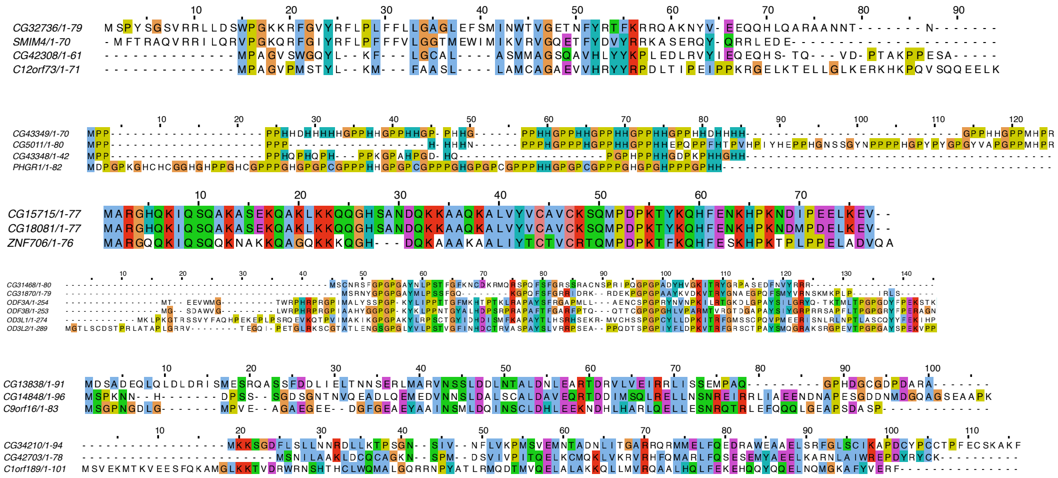

B

| Category | smORF gene | Human Homologues | Type of KO | # alleles | Phenotype |
|---|---|---|---|---|---|
| Single gene KO | CG32736 | SMIM4 | Frameshift indel | 1 | lethal |
| | CG42308 | C12orf73 | Frameshift indel | 1 | lethal |
| | bc10 | BLCAP | Knock-in gene disruption | 2 | viable, fertile |
| | CG18508 | C18orf32 | Frameshift indel and large deletion | 4 | viable, fertile |
| | CG33169 | SMCO4 | Frameshift indel | 1 | viable, fertile |
| | CG42455 | SMIM13 | Frameshift indel | 2 | viable, fertile |
| | CG15715 | ZNF706 | Frameshift indel | 2 | viable, fertile |
| | CG17931 | SERF1A,SERF2,SERF1B | Frameshift indel | 2 | viable, fertile |
| | CG18081 | ZNF706 | Frameshift indel | 1 | viable, fertile |
| | CG18336 | FAM166A | Frameshift indel | 1 | viable, fertile |
| | CG32448 | SMIM8 | Frameshift indel | 2 | viable, fertile |
| | CG4101 | MANBAL | Frameshift indel | 1 | viable, fertile |
| | CG14818 | C9orf16 | Frameshift indel | 2 | viable, fertile |
| | CG31468 | ODF3L1, ODF3L2 | Frameshift indel | 2 | viable, fertile |
| | CG31870 | ODF3L1, ODF3L2 | Frameshift indel | 2 | viable, fertile |
| | CG42371 | CEBPZOS | Frameshift indel | 2 | viable, fertile |
| | CG42381 | CMC4 | Frameshift indel | 2 | viable, fertile |
| | CG42518 | NCBP2-AS2 | Frameshift indel | 2 | viable, fertile |
| | CG42703 | C1orf189 | Frameshift indel | 2 | viable, fertile |
| | CG45122 | SMKR1 | Frameshift indel | 2 | viable, fertile |
| | CG5011 | PHGR1 | Frameshift indel | 2 | viable, fertile |
| | CG43349 | PHGR1 | Frameshift indel | 1 | viable, fertile |
| | CG13838 | C9orf16 | Large deletion | 3 | viable, fertile |
| | CG34210 | C1orf189 | Large deletion | 3 | viable, fertile |
| | CG34228 | SMIM12 | Large deletion | 3 | viable, fertile |
| Multi gene KO | CG32736/CG42308 | SMIM4/C12orf73 | Large deletion | 1 | lethal |
| | CG43349/CG5011/CG43348 | PHGR1 | Large deletion | 1 | viable, fertile |
| | CG15715/CG18081 | ZNF706 | Large deletion | 1 | viable, fertile |
| | CG31468/CG31870 | ODF3L1, ODF3L2 | Frameshift indels | 1 combo | viable, fertile |
| | CG13838/CG14818 | C9orf16 | Frameshift indel and Large deletion | 1 combo | viable, fertile |
| | CG34210/CG42703 | C1orf189 | Large deletion and frameshift indel | 1 combo | TBD |

C

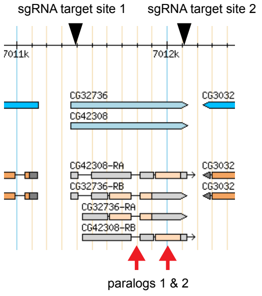

D

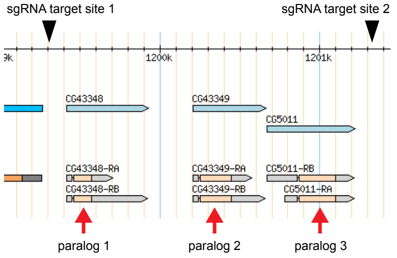

**Figure S6 – Full CRISPR knockout of 25 uncharacterized smORF genes and muli-gene knockout of paralogs. (*A*)** Protein sequence alignment of smORF paralogs listed in Figure 2B with their human homologs. **(B)** List of 25 uncharacterized smORFs, their human homologs and fly paralogs, type of molecular knockout, and homozygous fly phenotype. **(C, D)** Genomic views paralogs with location of sgRNA target sites for multi-paralog deletion.
