## Supplemental File 3 for "Molecular and functional characterization of the *Drosophila melanogaster* conserved smORFome"

| *Gene* | *cDNA* | *Max Expression/Stage, Condition* | *Max Score* | *Embryonic pattern* | *Spatial Expression Annotations** | *Organ System* |
|---|---|---|---|---|---|---|
| *Acbp1* | IP02950 | WPP_SalGlands | 454 | 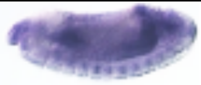 | midgut, hindgut, ubi | Endo/Midgut: HiGut |
| *Acbp2* | RH39533 | em10-12hr | 118 | 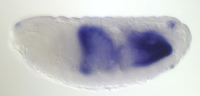 | midgut, fat body | Endo/Midgut: Blood/Fat |
| *Acbp4* | RE33457 | L2 | 332 | 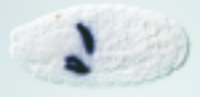 | gastric caecum | Endo/Midgut |
| *Acbp5* | RE05521 | AdMMF_Ecl_1d_DigSys | 2520 | 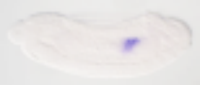 | midgut | Endo/Midgut |
| *Atox1* | LD15555 | WPP_24hr | 24.6 | 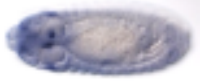 | crystal cell, ubi | Blood/Fat |
| *ATPsynE*[†] | GH08548 | WPP_4d | 275 | 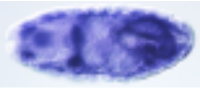 | DPPM, somatic muscle, midgut, MT, hindgut, ubi | Meso/Muscle: Endo/Midgut |
| *ATPsynεL*[†] | IP03962 | AdMM_Ecl_4d_Testes | 23.2 | 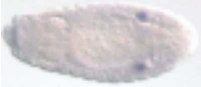 | gonad, faint ubi | Pole/Germ Cell |
| *ATPsynG*[†] | RE20862 | AdMMF_Ecl_1d_Carcass | 427 | 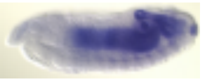 | DPPM, muscle, midgut, MT, hindgut, anal pad | Meso/Muscle: Endo/Midgut: HiGut |
| *Baf* | GH06291 | em4-6hr | 463 | 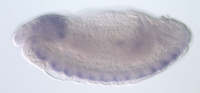 | brain, VNC, ubi | CNS |
| *BBIP1/ CG8135* | GH05505 | AdMM_Ecl_4d_Testes | 22.5 | 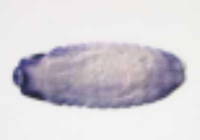 | clypeolabrum, V and head epid | FoGut:Ect/Epi |
| *bc10* | LD43519 | AdMF_Ecl_4d_Ovaries | 225 | 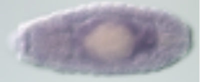 | midgut, MT, ubi | Endo/Midgut: HiGut |
| *bol* | GH12029 | em8-10hr | 27.6 | 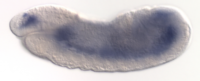 | Somatic muscle | Meso/Muscle |
| *Ccdc56*[†] | MIP03077 | AdMM_Ecl_4d_Testes | 115 | 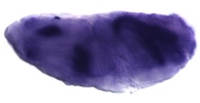 | DPPM, somatic muscle, midgut, MT, hindgut, ubi | Meso/Muscle: Endo/Midgut |
| *cer* | LP06209 | WPP Salivary Glands | 1220 | 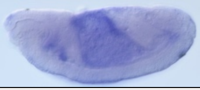 | amnioserosa | ExtraEmb |
| *CG11825*[†] | RE25483 | em2-4hr | 26.5 | 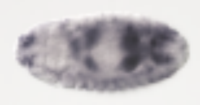 | clypeolabrum, D and V epid, midgut, MT, ubi | FoGut:Ect/Epi Endo/Midgut: HiGut |
| *CG12355*[†] | RE25221 | AdM_Ecl_1d | 4.27 | 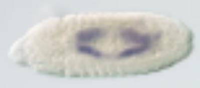 | midgut, MT | Endo/Midgut: HiGut |

| | | | | | | |
|---|---|---|---|---|---|---|
| *CG12384* | RH17411 | AdMMF_Ecl_1d_DigSys | 224 | 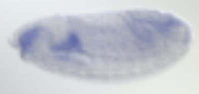 | D and V epi, SG, foregut, hindgut | Ect/Epi:SalGl: FoGut:HiGut |
| *CG13018*† | IP03340 | AdMM_Ecl_4d_Testes | 17.1 | 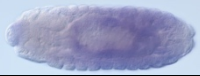 | midgut | Endo/Midgut |
| *CG13364* | LP11709 | AdMM_Ecl_4d_AccGl | 226 | 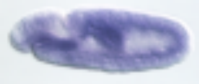 | midgut, muscle, ubi | Meso/Muscle: Endo/Midgut |
| *CG13784* | SD10385 | WPP Fat Body | 35.2 | 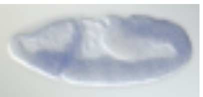 | ant endoderm, head and trunk mesoderm | Endo/Midgut: Meso/Muscle |
| *CG14036* | FI16614 | AdVF_Ecl_4d_Ovaries | 432 | 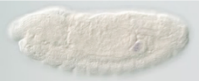 | gonad | Pole/Germ Cell |
| *CG14104* | IP05391 | L3_Carcass | 32.4 | 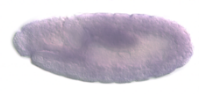 | midgut, ubi | Endo/Midgut |
| *CG14483*† | IP03474 | AdMM_Ecl_4d_Testes | 68.8 | 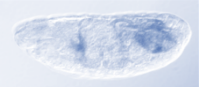 | midgut, faint ubi | Endo/Midgut |
| *CG14645* | GH16406 | AdMMF_Ecl_1day_DigSys | 2350 | 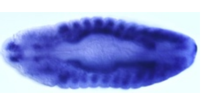 | DPPM, somatic muscle, visceral muscle | Meso/Muscle |
| *CG14933* | RH17612 | AdMM_Ecl_20d_Heads | 558 | 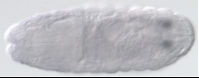 | posterior spiracle | Tracheal |
| *CG15386*† | LP21834 | L2 | 28.2 | 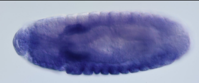 | salivary gland, ubi | SalGl |
| *CG15715* | MIP04739 | AdMF_Ecl_4d_Ovaries | 103 | 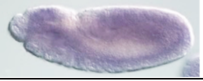 | head and trunk mesoderm, brain, ubi | Meso/Muscle: CNS |
| *CG17734*† | FI02890 | AdVF_Ecl_4d_Ovaries | 46.2 | 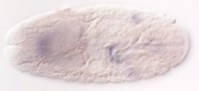 | gonad | Pole/Germ Cell |
| *CG17776* | IP03739 | AdMM_Ecl_4d_Testes | 65.2 | 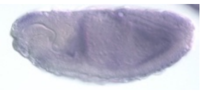 | DPPM, midgut, hindgut, faint ubi | Meso/Muscle: Endo/Midgut: HiGut |
| *CG18428* | MIP13681 | WPP_2d_CNS | 22.7 | 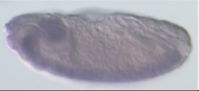 | brain, VNC | CNS |
| *CG30109* | AT19138 | AdMM_Ecl_4d_Testes | 49.9 | 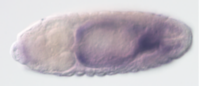 | midgut, hindgut | Endo/Midgut: Higut |
| *CG31262* | IP12186 | AdVF_Ecl_4d_Ovaries | 3.1 | 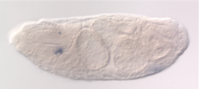 | crystal cell | Blood/Fat |
| *CG31313* | RH02566 | em10-12hr | 179 | 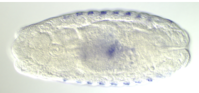 | Epidermis, yolk nuclei | Ect/Epi: Extraemb |
| *CG32069* | IP03767 | em10-12hr | 18.5 | 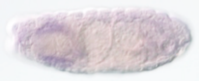 | SG, ubi | SalGl |

| | | | | | | |
|---|---|---|---|---|---|---|
| *CG32276* | RE27904 | WPP_FatBody | 231 | 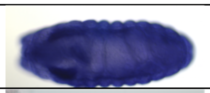 | D, V and head epid, SG, yolk nuclei, ubi | Ect/Epi:SalGl:Extraemb |
| *CG32448* | IP20115 | AdM_Ecl_30d | 137 | 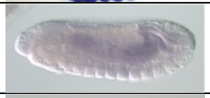 | midgut | Endo/Midgut |
| *CG33169* | RH56317 | AdMM_Ecl_4d_AccGl | 37.4 | 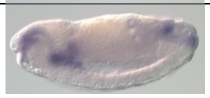 | foregut, SG, anal pad, post spiracle | FoGut:SalGl: Ect/Epi: Tracheal |
| *CG33170* | FI16530 | AdMM_Ecl_4d_Testes | 72.4 | 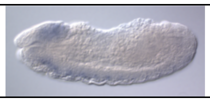 | brain, VNC, faint ubi | CNS |
| *CG33672/ Mvk* | MIP03836 | AdMF_Ecl_4d_Ovaries | 114 | 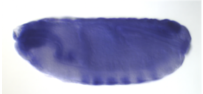 | D, V and head epid, foregut, midgut, MT, hindgut, muscle | Ect/Epi:FoGut: Endo/Midgut: HiGut:Meso/ Muscle |
| *CG33774* | GM20929 | em4-6hr | 33 | 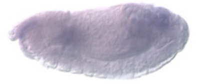 | midgut, ubi | Endo/Midgut |
| *CG34132*[†] | IP17640 | L2 | 64.9 | 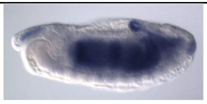 | DPPM, fat body, midgut, hindgut, anal pad, ubi | Meso/Muscle: Blood/Fat Endo/Midgut: Higut/Ect/Epi |
| *CG34242*[†] | IP17193 | AdMMF_Ecl_1d_Carcass | 35.8 | 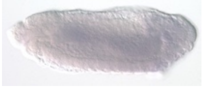 | midgut, ubi | Endo/Midgut |
| *CG34276* | BS27396 | L3_SalivaryGlands | 20.4 | 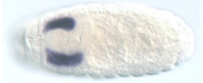 | SG | SalGl |
| *CG40472*[†] | PC01010 | WPP_3d | 51.8 | 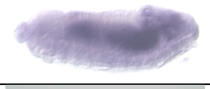 | midgut, midgut chamber, ubi | Endo/Midgut |
| *CG4101* | IP02903 | WPP_SalivaryGlands | 31.8 | 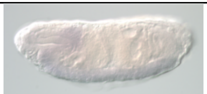 | brain, VNC | CNS |
| *CG41128*[†] | RH61753 | AdMF_Ecl_4d_Ovaries | 9.6 | 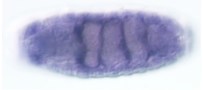 | midgut chamber, MT, ubi | Endo/Midgut: HiGut |
| *CG42259* | GH03322 | AdVF_Ecl_20d_Heads | 19.5 | 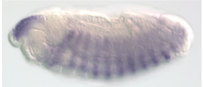 | DPPM, somatic muscle | Meso/Muscle |
| *CG42371/ CG15386* | GH19557 | L2 | 28.2 | 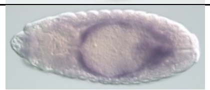 | DPPM, midgut, hindgut, ubi | Meso/Muscle Endo/Midgut: HiGut |
| *CG42375*[†]*/ CG9288* | LD32260 | em2-4hr | 31.4 | 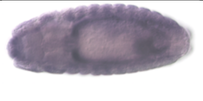 | DPPM, midgut, hindgut, ubi | Meso/Muscle: Endo/Midgut: HiGut |
| *CG42381*[†]*/ CG42379/ CG42380/ Pig-M* | GH02741 | AdMM_Ecl_4d_Testes | 37 | 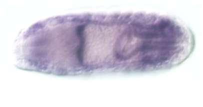 | epi- and hypo-pharynx, DPPM, foregut, midgut, MTT, hindgut, ubi | FoGut: Endo/Midgut: HiGut |

| | | | | | | |
|---|---|---|---|---|---|---|
| *CG42394* | MIP05983 | WPP_24hr | 18.5 | 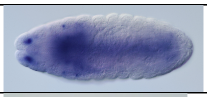 | brain, VNC | CNS: PNS |
| *CG42455/ vnc* | RE01326 | AdMM_Ecl_4d_Testes | 53.7 | 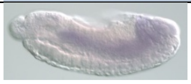 | midgut, ubi | Endo/Midgut |
| *CG42486* | GH09228 | AdMM_Ecl_20d_Heads | 558 | 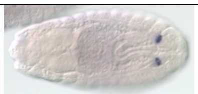 | posterior spiracle | Tracheal |
| *CG42496*[†]*/ Ppcdc* | LD37882 | AdMM_Ecl_4d_Testes | 63.1 | 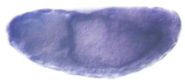 | midgut, ubi | Endo/Midgut |
| *CG42497*[†]*/ Tim10*[†] | LD46744 | AdMM_Ecl_4d_Testes | 242 | 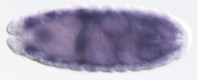 | DPPM, midgut, hindgut, MT, ubi | Meso/Muscle: Endo/Midgut; HiGut |
| *CG42587* | FI17589 | L3_FatBody | 582 | 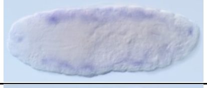 | Fat Body | Blood/Fat |
| *CG42740* | GH11688 | WPP_2d_CNS | 27.6 | 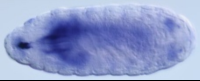 | brain, VNC | CNS |
| *CG43058* | PC01017 | AdMM_Ecl_4d_Testes | 44 | 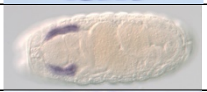 | SG | SalGl |
| *CG44008* | LP11341 | L3_DigSys | 533 | 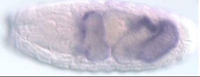 | midgut, midgut chamber | Endo/Midgut |
| *CG44194*[†]*/ CG17327* | MIP04754 | em0-2[‡] | 72 | 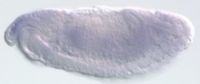 | DPPM, midgut, faint ubi | Meso/Muscle: Endo/Midgut |
| *CG5011* | LP08456 | AdMMF_Ecl_1d_DigSys | 461 | 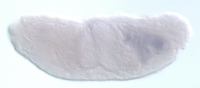 | midgut | Endo/Midgut |
| *CG5446* | RE36920 | WPP_FatBody | 165 | 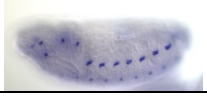 | sensory nervous sys, ubi | PNS |
| *CG6770* | RH17958 | WPP_SalGlands | 13500 | 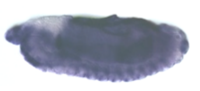 | amnioserosa, SG, crystal cell, vent midline, foregut, anal pad, ubi | Extraemb: SalGl: Blood/Fat:CNS :FoGut:Ect/Epi |
| *CG6878*[†] | IP03042 | AdVF_Ecl_4d_Ovaries | 115 | 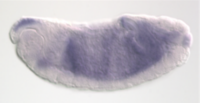 | midgut, muscle, faint ubi | Meso/Muscle: Endo/Midgut |
| *CG7637* | IP03042 | AdVF_Ecl_4d_Ovaries | 115 | 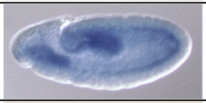 | midgut, muscle, faint ubi | Meso/Muscle: Endo/Midgut |
| *CG7646* | GM14157 | AdMMF_Ecl_4d_Carcass | 30.6 | 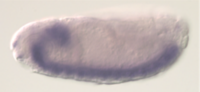 | brain, VNC | CNS |
| *CG8860* | GM14157 | AdMMF_Ecl_4d_Carcass | 383 | 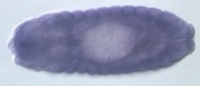 | proventriculus, muscle, midgut, ubi | FoGut:Meso/ Muscle: Endo/Midgut |

| | | | | | | |
|---|---|---|---|---|---|---|
| *cib* | RE37354 | em0-2hr | 2490 | 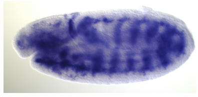 | brain, VNC, muscle, trachea, plasmatocytes, ubi | CNS: Meso/Muscle: Tracheal |
| *Cks30A* | LD20271 | AdVF_Ecl_4d_Ovaries | 78.5 | 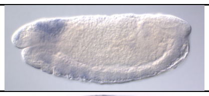 | brain, VNC, gonad | CNS: Pole Cells |
| *COX6B*[†] | GH28726 | AdMMF_Ecl_1d_Carcass | 721 | 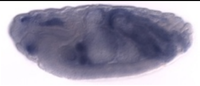 | DPPM, midgut, hindgut, ubi | Meso/Muscle: Endo/Midgut: HiGut |
| *COX7A*[†] | GM26747 | AdMMF_Ecl_4d_Carcass | 322 | 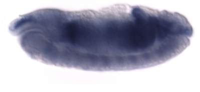 | DPPM, midgut, MT, hindgut, muscle, ubi | Meso/Muscle: Endo/Midgut: HiGut |
| *COX7C*[†] | LD14731 | AdMMF_Ecl_1d_Carcass | 602 | 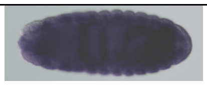 | DPPM, midgut, hindgut, muscle, ubi | Meso/Muscle: Endo/Midgut: HiGut |
| *COX8*[†] | GH15688 | WPP_4d | 391 | 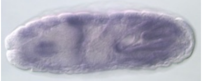 | DPPM, MT, midgut, hindgut, muscle, ubi | Meso/Muscle: Endo/Midgut: HiGut |
| *cpx* | MIP27931 | AdMM_Ecl_1d_Heads | 172 | 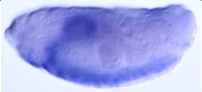 | brain, VNC | CNS |
| *ctp* | LD24056 | AdMMF_Ecl_4d_Carcass | 194 | 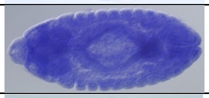 | brain, VNC, PNS, midgut, ubi | CNS: PNS: Endo/Midgut |
| *cype*[†] | GH04604 | AdMMF_Ecl_1d_Carcass | 450 | 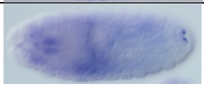 | DPPM, midgut, posterior spiracle, ubi | Meso/Muscle: Endo/Midgut: Trachae |
| *Dpm2* | SD05789 | WPP_FatBody | 304 | 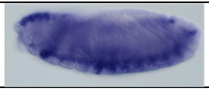 | DPPM, somatic muscle, ubi | Meso/Muscle |
| *drm* | LD26791 | em10-12hr | 20.7 | 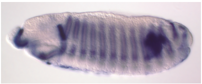 | Epidermis, MT, Trachea, Proventriculus, Lymph Gland, Pericardial cell | Tracheal |
| *EMRE*[†] | RE55001 | em2-4hr | 55.4 | 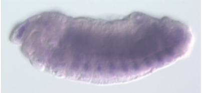 | DPPM, somatic muscle, ubi | Meso/Muscle |
| *Gγ30A* | MIP20711 | AdMM_Ecl_1d_Heads | 184 | 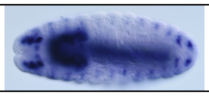 | brain, VNC, sensory system head | CNS/PNS |
| *glob1* | GH09296 | em10-12hr | 70.9 | 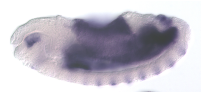 | fat body, amnioserosa, hypopharynx | Blood/Fat |
| *Ilk* | LD24671 | em10-12hr | 67.1 | 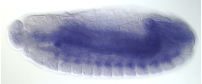 | muscle system, garland cell | Meso/Muscle Blood/Fat |

| | | | | | | |
|---|---|---|---|---|---|---|
| *IM33* | RH05411 | AdVF_Ecl_20d_Heads | 1100 | 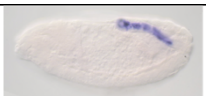 | large intestine, garland cells, fat body | HiGut: Blood/Fat |
| *IM33* | RH05411 | AdVF_Ecl_20d_Heads | 1100 | 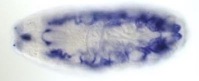 | large intestine, garland cells, fat body | HiGut: Blood/Fat |
| *ksh* | IP03267 | AdF_Ecl_5d | 28.6 | 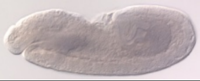 | midgut | Endo/Midgut |
| *kud* | MIP32344 | L2 | 45.1 | 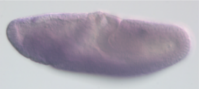 | SG, ubi | SalGl |
| *mbl*‡ | RH33825 | WPP-2d_CNS | 51 | 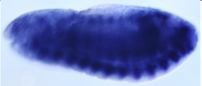 | muscle | Meso/Muscle |
| *Mlp60A* | RH74176 | em10-12hr | 451 | 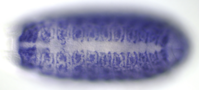 | DPPM. muscle sy, somatic muscle, visceral muscle, | Meso/Muscle |
| *mRpS21*† | RE23482 | L2 | 33.1 | 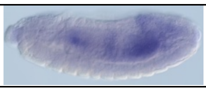 | DPPM, midgut, MT | Endo/Midgut: Meso/Muscle: |
| *MsrA* | FI09229 | em10-12hr | 375 | 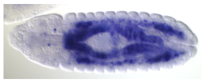 | fat body, midgut, epidermis, lymph gland, crystal cell, yolk nuclei | Blood/Fat |
| *ND-15*† | GH23780 | em10-12hr | 78.8 | 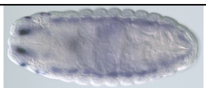 | DPPM, midgut, MT hindgut, | Meso/Muscle:Endo/Midgut:HiGu |
| *ND-AGGG*† | RH18150 | AdM_Ecl_1d | 79.4 | 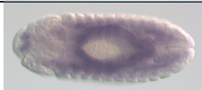 | DPPM, midgut, visceral muscle, faint ubi | Meso/Muscle: Endo/Midgut |
| *ND-MLRQ*† | RE56733 | AdM_Ecl_1d | 21.4 | 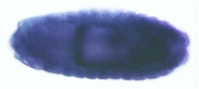 | DPPM, midgut, hindgut, muscle, strong ubi | Meso/Muscle: Endo/Midgut: HiGut |
| *ND-MNLL*† | GM22884 | WPP_4d | 194 | 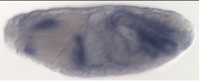 | DPPM, muscle, MT, midgut, hindgut, ubi | Meso/Muscle: Endo/Midgut: HiGut |
| *ND-MWFE*† | GH20835 | WPP_4d | 58 | 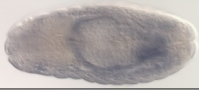 | midgut, faint ubi | Endo/Midgut |
| *Neb-cGP*† | RE31692 | WPP_4d | 302 | 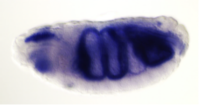 | DPPM, midgut, MT, hindgut, muscle, ubi | Meso/Muscle: Endo/Midgut: HiGut |
| *Nxt1* | RE28995 | em2-4hr | 475 | 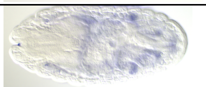 | gonad, ubi | Pole/Germ Cell |
| *osi*† | GH16758 | AdMMF_Ecl_4d_Carcass | 451 | 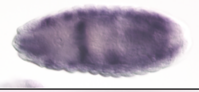 | DPPM, midgut, somatic muscle, ubi | Meso/Muscle: Endo/Midgut |
| *ox*† | GM22210 | WPP_4d | 365 | 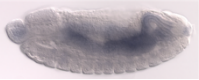 | midgut, ubi | Endo/Midgut |

| | | | | | | |
|---|---|---|---|---|---|---|
| *peg* | SD04019 | em10-12hr | 70.3 | 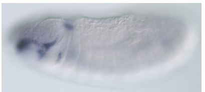 | maxillary, labial, dorsal/lateral sensory complexes, crystal cell | PNS Blood/Fat |
| *Pis* | RE35104 | em0-2hr | 55.2 | 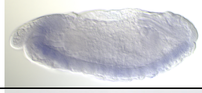 | brain, VNC, ubi | CNS |
| *Polr2L* | SD08670 | em4-6hr | 49.3 | 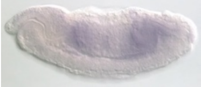 | midgut, MT, hindgut, fat body, muscle | Endo/Midgut: Meso/Muscle: Blood/Fat |
| *Polr2K* | IP17848 | AdVF_Ecl_4d_Ovaries | 42.4 | 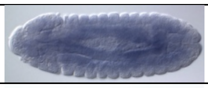 | midgut, muscle | Endo/Midgut: Meso/Muscle |
| *Pop5* | IP04375 | em4-6hr | 25.5 | 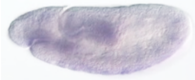 | ant and post midgut | Endo/Midgut |
| *REPTOR-BP* | RE65609 | L3_DigSys | 211 | 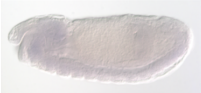 | brain, VNC, faint ubi | CNS |
| *RNASEK* | GM16138 | em22-24hr | 107 | 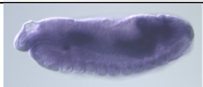 | midgut, MT, hindgut, ubi | Endo/Midgut: HiGut |
| *RpS28a* | RT02924 | AdMM_Ecl_4d_Testes | 50.8 | 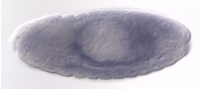 | midgut, ubi | Endo/Midgut |
| *Sdc* | LD08230 | em4-6hr | 26.5 | 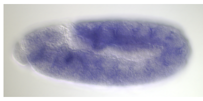 | brain, VNC, muscle, foregut, midgut, hindgut, SNS, epidermis | CNS |
| *Sec61β* | RH61539 | em10-12hr | 247 | 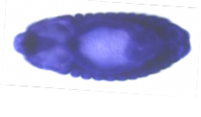 | atrium, D, V and head epid, SG, foregut, midgut, hindgut, ubi | Ect/Epi:SalGl: FoGut/Endo/ Midgut:Higut |
| *Sec61γ* | RE69515 | em10-12hr | 112 | 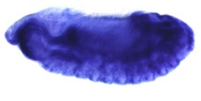 | foregut, proventriculus, hindgut, salivary gland | FoGut: HiGut: SG |
| *Sem1* | RH34416 | L3_Carcass | 374 | 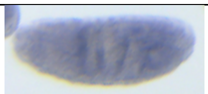 | midgut, ubi | Midgut |
| *Sf3b5* | IP03424 | em4-6hr | 105 | 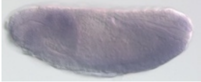 | brain, VNC, faint ubi | CNS |
| *SLIRP1* | GM17572 | em10-12hr | 241 | 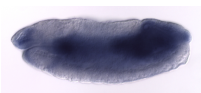 | midgut, hindgut, muscle | Endo/Midgut: Meso/Muscle: HiGut |
| *SLIRP2* | IP20549 | L1 | 13.7 | 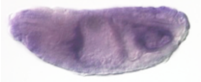 | midgut, ubi | Endo/Midgut |
| *sloth1*[†]/ *sloth2*[†] | RE60462 | WPP_3days | 19.5 | 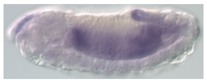 | DPPM, midgut, hindgut, MT | Meso/Muscle: Endo/Midgut: HiGut |

| | | | | | | |
|---|---|---|---|---|---|---|
| *SmE* | GM19936 | em4-6hr | 154 | 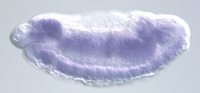 | brain, VNC, foregut, midgut, hindgut, ubi | CNS: FoGut: Midgut: HiGut |
| *SmF* | GM23968 | em2-4hr | 134 | 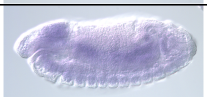 | brain, VNC, midgut, ubi | CNS: Midgut |
| *SNRPG* | RH35475 | AdVF_Ecl_4d_Ovaries | 120 | 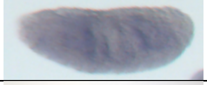 | midgut, ubi | Midgut |
| *Spase12* | RE02772 | em10-12hr | 300 | 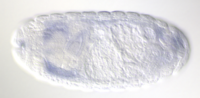 | SG | SalGl |
| *Spec2* | LD24607 | em2-4hr | 77.4 | 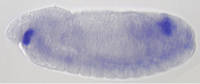 | hypopharynx, brain, VNC, posterior spiracle | FoGut: CNS: Tracheal |
| *Srp9* | RE23260 | em10-12hr | 215 | 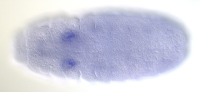 | SG, ubi | SalGl |
| *Sumo* | LD07775 | L3_CNS | 1270 | 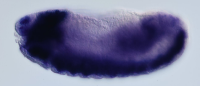 | brain, VNC, labral, dorsal/lateral sensory complexes | CNS: PNS |
| *sun*[†] | RE19513 | AdMMF_Ecl_1d_Carcass | 281 | 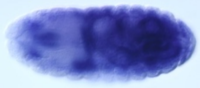 | DPPM, muscle, MT, midgut, hindgut, ubi | Meso/Muscle: Endo/Midgut: HiGut |
| *Svip* | GH02734 | WPP_SalGlands | 44.4 | 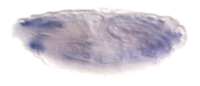 | DPPM, somatic muscle | Meso/Muscle |
| *Tango11* | LD25567 | AdMM_Ecl_4d_Testes | 70.7 | 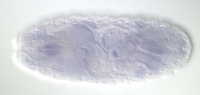 | gonad, ubi | Gonad |
| *Tfb5/ CG31917* | RE52596 | AdMM_Ecl_4d_Testes | 189 | 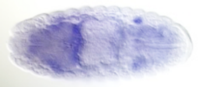 | midgut, gonad, faint ubi | Endo/Midgut |
| *Tim8*[†] | SD06593 | AdVF_Ecl_4d_Ovaries | 136 | 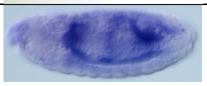 | DPPM, midgut, hindgut, somatic muscle, ubi | Meso/Muscle: Endo/Midgut: HiGut |
| *Tim9a*[†] | IP02031 | AdMM_ecl_4d_Testes | 36.5 | 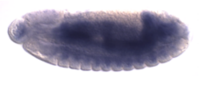 | DPPM, midgut, hindgut, visceral muscle, ubi | Meso/Muscle: Endo/Midgut: HiGut |
| *Tina-1* | GH28534 | em10-12hr | 375 | 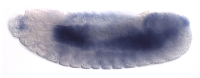 | DPPM, Midgut, ubi | Meso/Muscle: Endo/Midgut |
| *Tom7*[†] | AT23266 | L2 | 132 | 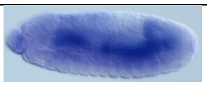 | DPPM, midgut, hindgut, viceral muscle, ubi | Meso/Muscle: Endo/Midgut: HiGut |
| *Ufm1* | MIP14123 | AdMM_Ecl_4d_AccGl | 46.9 | 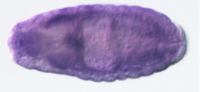 | clypeolabrum, epi- and hypo-pharynx, SG, trachea, ubi | FoGut: SalGl: Tracheal |

| | | | | | | |
|---|---|---|---|---|---|---|
| *UQCR-6.4*† | RH56961 | WPP_4d | 195 | 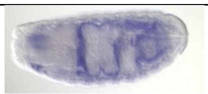 | DPPM, midgut, MT, hindgut, muscle, faint ubi | Meso/Muscle: Endo/Midgut: HiGut |
| *UQCR-11*† | RH21839 | AdMMF_Ecl_1d_Carcass | 106 | 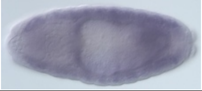 | muscle, MT, midgut, ubi | Meso/Muscle: Endo/Midgut: |
| *UQCR-Q*† | AT13736 | AdMMF_Ecl_1d_Carcass | 506 | 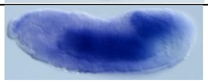 | DPPM, midgut, hindgut, muscle, ubi | Meso/Muscle: Endo/Midgut: HiGut |
| *VhaM9.7-a* | IP04021 | em10-12hr | 41 | 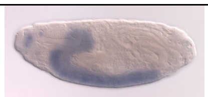 | brain, VNC, Sensory System Head | CNS |
| *VhaM9.7-b* | FI02825 | em0-2hr | 343 | 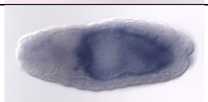 | midgut | Endo/Midgut |
| *Vti1b*‡ | FI17308 | WPP_FatBody | 22 | 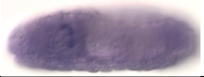 | Garland cell | Meso/Midgut |

**Abbreviations: AccGl-Accessory Gland;* AdMMF-*Adult Mixed Male Female; ant-anterior; D-dorsal; DPPM-dorsal prothoracic pharyngeal muscle; Ecl-Eclosed; epid-epidermis; MT-Malpighian tubules; post-posterior; SG-salivary gland; ubi-ubiquitous; V-ventral; VNC- Ventral nerve cord; WPP-White Pre-Pupae*

*†Mitochondrially expressed*

*‡Max Expression Stage and Score from FlyBase, all others from modENCODE (Graveley et al., 2011)*
