## Supplemental File 6 for "Molecular and functional characterization of the *Drosophila melanogaster* conserved smORFome"

### **Materials**

**Reagents (In order of first appearance in protocol)**

- *Drosophila melanogaster,* OregonR strain
- Water (deionized)
- Sucrose (Sigma, cat. no. S0389)
- Ethanol (Sigma, cat. no. E7023)
- Tris-Cl (20 mM, pH7.4, cat. no.)
  - 1M Tris pH 7.0 (Invitrogen, cat. no. AM9850G)
  - 1M Tris pH 8.0 (Invitrogen, cat. no. AM9855G)
- KCl (140 mM, Invitrogen, cat. no. AM9640G)
- MgCl_2_ (5 mM, Invitrogen, cat. no. AM9530G)
- DTT (0.5 mM, Sigma, cat. no. 43816)
- Tergitol Solution (1% (V/V), Sigma, Cat. No. NP40S)
- RNasin Plus RNAse Inhibitor (200 U/ 1μg, Promega, cat. No. N2111 or N2115, depending on number of samples)
- Turbo DNaseI (Invitrogen, cat. No. AM2239)
- 5.7% Clorox bleach
- Harringtonine (LKT Laboratories, cat. no. H0169)
- Cycloheximide (100 μg/ml) (Sigma Aldrich, cat. no. C4859-1ML)
- TRIzol Reagent (Thermo Fisher, cat. no. 15596026)
- 3 M NaOAc pH5.2 (Sigma Aldrich, Cat. no. S7899)
- GlycoBlue (Thermo Fisher, cat. no. AM9515)
- RNaseI (100U/μl) (Ambion, cat. no. AM2294)
- Micro RNA Gel Ladder (NEB, Cat. No. N2102S)
- Low Range ssRNA Ladder (NEB, Cat. No. N0364S)
- 31 nt Oligonucleotide Size Marker (Upper Marker) (IDT, Custom Order)

5’-rArUrGrUrArCrArCrGrGrArGrUrCrGrArCrCrUrCrArArCrCrCrGrCrArArCrGrCrGrA/3Phos-3’

- 26 nt Oligonucleotide Size Marker (Lower Marker) (IDT, Custom Order)

5’-rArUrGrUrArCrArCrGrGrArGrUrCrGrArCrCrCrArArCrGrCrGrA/3Phos-3’

- SYBR Gold nucleic acid gel stain (Invitrogen by Thermo Fisher Cat. No.
- Monarch PCR and DNA Kit (New England Bio (NEB), cat. no. T1030)
- Novex TBE-Urea Gels, 15%, 12 well (Thermo Fisher, cat. no. EC68852BOX)
- Novex TBE-Urea Sample Buffer (2X) (Thermo Fisher, cat. no. LC6876)
- Novex TBE-Running Buffer (5X) (Thermo Fisher, cat. no. LC6675)
- Zymo Research Small–RNA PAGE Recovery Kit (Zymo Research, cat. no. R1070)
- NEBNext Small RNA Library Prep Set for Illumina or NEBNext Multiplex Small RNA library Prep Set for Illumina (New England Bio (NEB), Catalog number dependent on the number of libraries being processed for whole-genome sequencing)

**Optional Reagents**

- Gel Set-up
  - microRNA oligonucleotide size markers
  - 28 bp and 30 bp oligonucleotide size markers

**Equipment**

- Fly cages
- 1.5 ml Microtubes (Axygen, MCT-150-C)
- Pellet pestles (Sigma, cat. no. Z359947)
- Dounce homogenizer Corning Pyrex 7727-2 Ten Broeck Homogenizer with Pour Spout, 2 Ml
- Beckman Ultra-clear Centrifuge Tubes, 14 x 89 mm, (Beckman Coulter, cat. no. 344059)
- Axygen Microtubes (MCT-150-C)
- Molasses plates (Preparation method detailed below)
- Hoefer Gradient Maker (SG30)
- Thermocycler (Perkin Elmer, GeneAmp PCR System 9700, Part no. N8050200)
- Hypodermic needle (22G x 4, Air-Tite, cat. no. 8300014471)
- MasterFlex Peristaltic Pump C/L Drive (Cole Parmer, cat. No. EW77120-62 or 77122-14)
- Beckman Coulter Ultracentrifuge (Model Optima L-90)
- Beckman Coulter Ultracentrifuge Rotor (cat. no. SW41ITI)
- NanoDrop ND-1000 Spectrofluorometer (cat. no. E112352)
- Qubit 3.0 Fluorometer (cat. no. Q33216)
- Invitrogen XCell SureLock Minicell (Invitrogen, cat. No. 1188039-2704)
- dounce homogenizer

**Reagent Setup and Preparation**

- Preparation of 10% and 50% w/v sucrose solutions - TIMING 15 min. per sample
  a. 20 mM Tris-Cl (pH7.4), 140 mM KCl, 5 mM MgCl2,0.5 mM DTT, 1% (V/V) Triton-X,100 μg/mL cycloheximide, 200U/1ug RNasin Plus RNAse Inhibitor, 25U/ml Turbo DNaseI
- Molasses food tray preparation
- **Reproductive toxin preparation**
  a. Harringtonine Prep
  b. Cycloheximide Prep
- **Prepare 10%-50% sucrose gradient (15’ tube)**

Beckman 344059 (14 x 89 mm) Ultra-clear centrifuge tubes and Hoefer Gradient maker

a. Pre-Chill UltraCentrifuge Rotor at 4°C
b. Pour gradient from TOP
   i. Use MasterFlex C/L peristaltic pump
   ii. Use Long Hypodermic needle (22G x 4) (Air-Tite 8300014471)
   iii. Tilt the ultracentrifuge tube (rest it on the 50ml falcon tube rack)
   iv. Hang the Long Hypodermic needle attached to tubing from a stand so that it just reaches inside the tilted tube
   v. Place identical stir bars to each chamber of SG30 gradient maker
c. 15mm max length
d. Confirm that both stopcocks are closed
   i. Add 5.6 ml of 10% sucrose to the reservoir chamber (back) of the gradient maker
e. Open connector stopcock and allow the solution to flow through connector channel to the edge of the mixing chamber, then close the stockcock
f. Be sure no large bubbles obstruct the flow
   i. Add 5.6 ml of 50% sucrose to the mixing chamber (front) of the gradient maker
g. Open Connector stopcock
h. Be sure no large bubbles obstruct the flow
   i. Store on ice until needed
i. Prepare identical sucrose gradient for the remaining samples
j. Alternatively make sure there is a balance tube for the rotor

### **Procedure**

1. ***Drosophila* Embryo Collections - TIMING 10 min. per tray**

Collect appropriately aged FRESH ModENCODE OregonR *D. Melanogaster* Embryos from [0–2, 2–4, 4–6, 10–12, 14–16 and 16–18hr] (10’ collection tray)

- Set up two houses with at least 40 bottles per house (14 g of flies)
- Collection in molasses trays
  1.1. Scrape yeast from agar plate if not too many embryos in the yeast
  1.2. Add deionized water to a tray and loosen embryos with a paint brush
  1.3. Pour embryos from the meat tray into a large three piece sieve and wash well to remove debris
  1.4. Transfer embryos to a small sieve and keep submerged in deionized water until ready to process
  1.5. De-chorionate embryos in sodium hypochlorite solution (5.7% Clorox bleach)
       1.5.1. Pour the hypochlorite solution into a large weight boat to cover embryos
       1.5.2. Incubate at RT for 3’ with gentle agitation
       1.5.3. Wash well with deionized water
       1.5.4. Dry embryos well
              1.5.4.1. Transfer most to previously weighed Eppendorf tube for Harringtonine processing
                       1.5.4.1.1. Weigh an Eppendorf Tube after adding the embryos
              1.5.4.2. Transfer some to another previously weighed Eppendorf tube for and save for RNA seq
                       1.5.4.2.1. Weigh an Eppendorf Tube after adding the embryos
                       1.5.4.2.2. Freeze in LN2 and place in –80 °C
  1.6. To collect two biological replicates for sequencing we started two fly cages - each founded with 14 g of adult flies (~14,000 adult OregonR flies). Cages were started 4 days before embryos were harvested, to acclimate the populations to the cage environment.
  1.7. Cages populations were maintained in Darwin Chamber Incubator under constant conditions of 27℃ and 70% humidity.
  1.8. Embryos were collected from molasses trays placed on the bottom of the fly cages.
  1.9. We sampled six embryonic stages across the 24 hours of *Drosophila* embryogenesis concurrently, and on the same day (2–4 hr., 4–6 hr., 6–8 hr., 12–14 hr., 16–18 hr., and 18–20 hr. aged embryos). All embryonic samples were immediately subjected to the ribosome profiling protocol (detailed below).
  1.10. For each 2 hr. developmental stage, a fraction of embryos were concurrently `flash frozen for paired RNA-seq analysis
  1.11. Non-embryonic tissues

2. **Harringtonine Pre-treatment and Embryo Homogenization - Timing 1 hr.**

- CAUTION! Harringtonine is highly toxic
- Dissolve 5 mg of Harringtonine powder in 500 ul of DMSO (to 10mg/ml); dispense into aliquots and store in the dark at –20 °C
- Dilute Harringtonine to a working concentration 2μg/ml (3μg/ml) in Mild Lysis Buffer w/o Cycloheximide
- Place diluted Harringtonine (3μg/ml) in Mild Lysis Buffer w/o Cycloheximide into 37°C
- Mild Lysis Buffer
  - 20 mM Tris-Cl (pH7.4), 140 mM KCl, 5 mM MgCl2,0.5 mM DTT, 1% (V/V) Triton-X, 0–100–200 μg/mL cycloheximide, 200U/1ug RNasin Plus RNAse Inhibitor, 25U/ml Turbo DNaseI
  2.1. Add 500μL of 2 μg/ml (3 μg/ml) Harringtonine (in MLB) to the embryos
  2.2. Incubate for 2’ at 37 °C *without homogenizing*
  2.3. Add 500μL ice–cold Cycloheximide (200 μg/ml) in MLB and mix well
  2.4. Homogenize w/ Blue Pestle for 20–30”
  2.5. Transfer to dounce homogenizer
  2.6. Stroke 15 x
  2.7. Add 500μL ice-cold Cycloheximide (100 μg/ml) in MLB and mix well
  2.8. Incubate on ice for 15’
  2.9. Centrifuge for 5’ at 8600 x g at 4 °C
  2.10. Transfer supernatant into clean microcentrifuge tube
- Avoid lipid layer and pellet
  2.11. Centrifuge for 5’ at 13000 x g at 4 °C
  2.12. Transfer supernatant into clean microcentrifuge tube
  2.13. Centrifuge for 10’ at 20200 x g at 4 °C
  2.14. Transfer supernatant into clean microcentrifuge tube (~1350 μl)
- Split Lysate into 3 aliquots and store at –80 °C
  - Aliquot 1 (450 μl) load onto sucrose gradient
  - Aliquot 2 (450 μl) Store at –80 °C
  - Aliquot 3 (~450 μl) Store at –80 °C
- ***SAFE STOPPING POINT Samples can be stored at –80 °C***

3. **Polysome Footprinting and precipitation**
   3.1. **Sucrose gradient and fractions collection - TIMING 3 hr. 30 min.**
        3.1.1. Load Aliquot 1 (450 μl) from **Step 3.14** onto sucrose gradient/s from **Step 1.4**
        3.1.2. Place tube w/ lysate on top of the gradient into Ultracentrifuge rotor
        3.1.3. Centrifuge for 2:30 at 36K at 4 °C
        3.1.4. Remove tubes from rotor buckets

- Use tweezers
- Seal with parafilm
- Attach working tube to clamps
- Keep remainder of the samples on ice
  3.1.5. Carefully pierce the bottom of the tube w/ 20 gauge needle attached to a syringe

and peristaltic pump

- Pierce the bottom of the tube just off center (to avoid the seam) at ~45° Collect ~24 drop fractions
- Keep collection plate on ice (to prevent RNA degradation)
- 1 drop= ~16.5 μL
- ~33 fractions
  3.1.6. Measure the absorbance of each fraction (A254)
- Use Plate Reader
  3.1.7. Seal plate and Freeze fractions at -80 °C until further use
- ***SAFE STOPPING POINT Samples can be stored at –80℃***
- Expecting ~12–16 polysomal fractions
  3.1.7.1. Combine all polysomal fractions and mix well
  3.1.7.2. Use 2ml to overlay over Sucrose Cushion Use 2ml to overlay over Sucrose Cushion 1 in **Step 4.1.7.4**
- Remainder is stored at –80 °C
  3.1.7.3. Add 10 ml of 34% Sucrose Cushion to an Ultrafuge tube
  3.1.7.4. Overlay mixed polysomal fractions [(~1.5ml)] over the Sucrose cushion
  3.2. **Sucrose Cushion (34%) #1 and RNaseI Digestion - TIMING 4 hr. 45 min. + Overnight**
       3.2.1. Centrifuge for 3:15 at 39K at 4 °C’
- Towards the end of the spin prepare 2ml Polysome Digestion Buffer/sample
- Use 15ul RNaseI (Ambion AM2294) (100U/μl) per 1 ml
- Label Eppendorf Tubes
  3.2.2. At the end of the spin pour off supernatant (sucrose)
  3.2.3. Fill ultrafuge tube with ice cold RNase Free water and immediately pour down to remove all sucrose (2x)
  3.2.4. Dry the pellets for 1’
- Usually no pellet is visible
  3.2.5. Add 875 μl of Polysome Digestion Buffer to the Ultrafuge tube seal with parafilm and mix well by vortexing
  3.2.6. Transfer the Polysome Digestion Buffer (w/ Polysomes) into a prelabeled Eppendorf tube
  3.2.7. Add additional 875 μll of Polysome Digestion Buffer into each sample
  3.2.8. Incubate for 45’ @ RT w/ gentle agitation
  3.2.9. Add 4.5 μl of SUPERase-In RNase Inhibitor to stop the RNAse I digestion
  3.3. **Sucrose Cushion (34%) #2 and RNA Isolation - TIMING 4 hr. 45 min. + Overnight + 45 min.**
       3.3.1. Add 9.75 ml of 30% Sucrose to an Ultrafuge tube
       3.3.2. Overlay digested polysomal fractions (~1.75ml) over the Sucrose cushion #2
       3.3.3. Centrifuge for 3hr 30 min at 39K at 4 °C
       3.3.4. Remove supernatant
       3.3.5. (2x) Fill tube with ice cold RNase Free Water and immediately pour down to remove all sucrose
       3.3.6. Dry the pellets for 1’
       3.3.7. Resuspend the pellet in 500 μl TRIzol
       3.3.8. Seal well w/ parafilm
       3.3.9. Vortex well
- Transfer to an Eppendorf Tube
  3.3.10. Add 0.1335 ml of Chloroform
  3.3.11. Vortex well
  3.3.12. Incubate at RT for 2’
  3.3.13. Spin for 5’ at 12000xg at 4 °C
  3.3.14. Transfer Aqueous (Usually upper) phase to a new pre-chilled Eppendorf tube
- ~250ul
  3.3.15. Add 28.1 μl 3M NaOAc- pH5.2, 4.2 μl GlycoBlue, 422 μl ice–cold Isopropanol
  3.3.16. Precipitate at –20 °C ON
- Or –80 °C for a minimum of 30’
  3.3.17. Spin at 4 °C at 20,800 *x g* for 30 min.
  3.3.18. Decant supernatant carefully and then spin briefly to collect the supernatant at the bottom of the tube. Remove the residual liquid with the micropipette.
  3.3.19. Air-dry the pellet for 1 min at room temperature.
  3.3.20. Resuspend in 10 μl of 10mM Tris–HCl pH8 and transfer to a clean Eppendorf Tube
- ***SAFE STOPPING POINT***
  3.4. **PAGE–UREA gel and Gel Extraction - TIMING 4 hr. 30 min.**
       3.4.1. To 10 μl of sample add 5 μl Novex TBE-Urea Sample Buffer (2x) [Invitrogen LC6876]
- Add 1 ul of 10 μM 34 nts upper oligo to 4 ul H20 and 5μl of Novex TBE-Urea Sample Buffer (2x)
- Add 1 ul of 10 μM 26 nts Lower oligo to 4 ul H20 and 5μl of Novex TBE-Urea Sample Buffer (2x)
- Add 1 ul of Low Range Marker to 4 ul H20 and 5μl of Novex TBE-Urea Sample Buffer (2x)
- Add 7.5 ul of microRNA marker (orange)
- Load every other well when loading to avoid contamination
  - Load buffer in all the wells regardless of the number of samples
- Load 26 nt lower oligo and 34 nt upper oligonucleotide markers
  3.4.2. Denature at 80°C for 90”
- XCell SureLock Minicell
  - Follow manufacturer’s instructions (Invitrogen) for gel box assembly, running of the gel and gel disassembly
- Pre–run gel for 15’
  3.4.3. Run 15% PAGE–UREA gel in 1xTBE for 65’ at 160V
- XCell SureLock Minicell
  - Follow manufacturer’s instructions (Invitrogen) for gel box disassembly
- Dilute SYBR Gold 1:10000 in TBE running buffer prior to use to obtain 1x solution
  3.4.4. Post–stain w/ SYBR Gold for 5’ at RT w/ gentle agitation
  3.4.5. Immediately excise gel slices
  3.4.6. Place into pre–labeled Epperdorf tubes
- Recover RNA from the PAGE–UREA Gel Slice by using slightly modified ZYMO RESEARCH ZR small–RNA™ PAGE Recovery Kit (R1070) protocol
- ~60–75”
  3.4.7. Crush the gel slice with a Zymo Research Squisher™-Single against the side of the Eppendorf tube. Add 400 µl RNA Recovery Buffer and incubate for 15’ at 65 °C (or until the gel slice dissolves)
- After initially crushing the gel slice against the side of the Eppendorf tube use pellet pestle motor for 20–30”
- Leave the Zymo Research Squisher™-Single in the tube while the tube is 65 °C and homogenize one more time for 20–30” after ~10’
  3.4.8. Transfer dissolved gel slices in RNA Recovery Buffer to Zymo-Spin™ IV Column (old FilterZymo-Spin™ III-F Filter) in a Collection Tube and Incubate for additional 5’
  3.4.9. Quick freeze the samples on dry ice or in a -80ºC freezer for 5 minutes, then transfer columns back into 65°C for 5 minutes to thaw.
  3.4.10. Snap theZymo-Spin™ IV Column tip and place the column back into a Collection Tube. Centrifuge at ≥1,500 × g for 30 seconds. Save the flow-through.
  3.4.11. Transfer the flow-through from the Step 4 to a Zymo-Spin™ IIICG Column in a Collection Tube and centrifuge at ≥1,500 × g for 30 seconds. Save the flow-through.
  3.4.12. Add 2 volumes of RNA MAX Buffer to the flow-through from Step 5 and mix well.
  3.4.13. Transfer the mixture to a Zymo-Spin™ IC Column in a Collection Tube. Centrifuge at ≥12,000 × g for 30 seconds. Discard the flow-through and place the Zymo-Spin™ IC Column back into the Collection Tube.
  3.4.14. Add 400 µl RNA Prep Buffer to the column. Centrifuge at ≥12,000 × g for 1 minute. Discard the flow-through and place the Zymo-Spin™ IC Column back into the Collection Tube.
  3.4.15. Add 800 µl RNA Wash Buffer to the column. Centrifuge at ≥12,000 × g for 30 seconds. Discard the flow-through and place the Zymo-Spin™ IC Column back into the Collection Tube.
  3.4.16. Repeat Step 9 with 400 µl RNA Wash Buffer.
  3.4.17. Centrifuge the Zymo-Spin™ IC Column at ≥12,000 × g for 2 minutes in an empty Collection Tube to ensure complete removal of the wash buffer.
  3.4.18. Place the Zymo-Spin™ IC Column into a provided DNase/RNase-Free Tube. Add 10 µl of the provided DNase/RNase-Free Water directly to the column matrix and let stand at room temperature for 1 minute.
  3.4.19. Centrifuge the Zymo-Spin™ IC Column at 10,000 × g for 1 minute to elute RNA. Recovered RNA can be used immediately or stored at ≤-70 °C
- ***PROCEED Directly to* Dephosphorylation**
  3.5. **End Repair (Dephosphorylation) - TIMING 2 hr. + Overnight**
- This dephosphorylation step is necessary to allow ligation of the 26-34 nt RNA fragments to the Universal miRNA linker.
  3.5.1. Add 33 μl of ice-cold MilliQ water to the 10 μl RNA
  3.5.2. Denature for 90 sec at 70 °C and then equilibrate at 37 °C.
  3.5.3. Add 5 μl of 10x polynucleotide kinase buffer, 1 μl of SUPERase-In and 1 μl of T4 polynucleotide kinase and mix well by pipetting.
  3.5.4. Incubate for 1 h at 37 °C and heat-inactivate for 10 min at 70 °C.
  3.5.5. Precipitate RNA by adding 39 μl of water, 1 μl of GlycoBlue, 10 μl of 3 M NaOAc-, mix well by inverting the tube, and then add 150 μl of ice-cold isopropanol. Mix well by inverting the tube. Incubate for at least 30 min at –80 °C (or on dry ice).
  3.5.6. Spin at 4°C at 20,800 *x g* for 30 min.
  3.5.7. Decant supernatant carefully and then spin briefly to collect the supernatant at the bottom of the tube. Remove the residual liquid with the micropipette.
  3.5.8. Air-dry the pellet for 1 min at room temperature.
  3.5.9. Re-suspend in 7 μl of 10mM Tris–HCl pH8 and transfer to a clean Eppendorf Tube
- *Measure OD*
  3.6. **NEBNext Small RNA Library Preparation - TIMING 5 hr.**
- NEBNext Small RNA Library Prep Set for Illumina was used to prepare NGS libraries with the standard NEB protocol, but with the following modifications:
  - Ligase concentration -NOT ligase, but Ligase adaptors and primers
  - 3’ SR Adaptor dilution 3:4 (1 ul of adaptor ; 1.33 nuclease-free water)
  - SR RT Primer 7:8 (1 ul of SR RT primer : 1.14 nuclease-free water)
  - 5’ SR Adaptor dilution 3:4 (1 ul of adaptor ; 1.33 nuclease-free water)
- Measure OD (NanoDrop)
  3.7. **Monarch PCR and DNA CleanUp Kit (#T1030)**
       3.7.1.1. To 100 μl of smRNA library add 700 μl Binding Buffer
- DO NOT Vortex; Mix by pipetting
  3.7.1.2. Add to Column
  3.7.1.3. Spin for 1’ at 12000xg at RT and discard flowthrough
  3.7.1.4. Re–insert the column and add 200 μl Wash Buffer
  3.7.1.5. Spin for 1’ at 12000xg at RT and discard flowthrough
  3.7.1.6. Re–insert the column and add another 200 μl Wash Buffer
  3.7.1.7. Spin for 1’ at 12000xg at RT and discard flowthrough
  3.7.1.8. Spin for 1’ at 12000xg at RT
- To assure that the tip of the column does not have any residual wash buffer
  3.7.1.9. Reinsert the column and elute w/ 27.5 μl of NC H20
  3.7.1.10. Spin for 1’ at 12000xg at RT
  3.7.2. Measure Concentration with Nanodrop and Quibit
- 1 μl to nanoDrop
- 1 μl to Qubit
- 1 μl to BioAnalyzer High-sensitivity DNA Chip
- ∼24 μl left
  3.7.3. Load onto Bionalzyzer (high sensitivity DNA chip)
  3.7.4. Normalize to 10 μM for NovaSeq
